# TRAMFIX: TRavelling Across Melbourne for FIXel-based analysis (a reproducibility study)

**DOI:** 10.1101/2025.07.21.666058

**Authors:** Remika Mito, Sila Genc, Jocelyn Halim, Joseph Yuan-Mou Yang, Jacques-Donald Tournier, Michael Kean, Chris Kokkinos, Richard McIntyre, Maria A. Di Biase, Robert Smith, Andrew Zalesky

## Abstract

Fixel-based analysis (FBA) has gained substantial interest for its ability to probe fibre-specific changes in the brain’s white matter from diffusion-weighted imaging data. However, the reproducibility and reliability of fixel-based measures across different scanners remains largely unknown. In this work, we present TRAMFIX: a multisite dataset of travelling participants (n=10 healthy adults) scanned across four 3T MRI scanners using a harmonised multi-shell DWI protocol. DWI data were processed using two pipelines that can adopted when performing multi-site FBA studies (site-specific versus pooled processing). We extracted fixel-based measures of fibre density (FD), fibre cross-section (FC), and fibre density and cross-section (FDC), as well as diffusion tensor imaging (DTI)-based fractional anisotropy (FA) and mean diffusivity (MD) from the harmonised protocol. Within-subject coefficients of variation (CV_ws_) and intraclass correlation coefficients (ICC) of FBA and DTI measures were computed at multiple resolutions of computation and analysis: (i) the whole-brain averaged level, (ii) tract-level, and (iii) fixel- or voxel-level. Fixel-based metrics demonstrated high reproducibility and reliability at the whole-brain level (CV_ws_ ranging between 0.51% to 1.57% and ICC between 0.93 and 0.996). While reproducibility and reliability remained high for tract-averaged FBA measures (particularly the FC and FDC metrics), some tracts exhibited lower ICC values < 0.8 for the FD measure. When examining fixel-level reliability and reproducibility clear spatial patterns emerged, with lower ICC across subcortical and cerebellar regions, and higher CV_ws_ at the cortical boundaries. Across all levels, tensor-based metrics demonstrated slightly lower reproducibility and reliability metrics than FBA measures, suggesting that FBA metrics are just as, if not more, reproducible than DTI metrics across scanners when using identical protocols. Our findings provide support for the reproducibility and reliability of fixel-based measures, highlighting their potential for use in multi-site FBA studies. Future work examining protocol-related differences, as well as appropriate harmonisation strategies when pooling data across sites and scanners, will be valuable.

## 1. Introduction

Diffusion-weighted imaging (DWI; or diffusion MRI) is currently the only tool available to non-invasively map the human brain’s complex white matter architecture. While the development of diffusion tensor imaging (DTI) was a revolutionary step in our ability to quantify changes to the brain’s white matter connections (Basser et al., 1994), known limitations to the technique’s modelling assumptions and biological interpretations (Jones et al., 2013) have led to major improvements in DWI acquisition and modelling approaches. The past few decades have seen a rapid rise in the development and application of these advanced DWI techniques, enabling improved characterisation of the brain’s complex white matter microstructure.

Fixel-Based Analysis (FBA; (Raffelt et al., 2017)) is one such framework that can provide more specific insight into the brain’s white matter fibre pathways. By modelling high-angular resolution diffusion imaging (HARDI) data with constrained spherical deconvolution (CSD; (Dell’Acqua & Tournier, 2019)), FBA can derive quantitative measures even in the presence of multiple fibre directions within voxels (known as ‘fixels’), enabling fibre-specific analysis of white matter pathways. The FBA framework has now been widely adopted across a range of clinical applications (see Dhollander et al., (2021) for a review), including in neurodegeneration (Ahmadi et al., 2024; Mito et al., 2018), psychopathology (Grazioplene et al., 2022; Kristensen et al., 2023, 2024; Lyon et al., 2019) and neurological disorders (Mito et al., 2022, 2024; Raffelt et al., 2017; Vaughan et al., 2017), as well as in healthy development (Genc et al., 2018; Genc, Malpas, et al., 2020) and aging (Kelley et al., 2021; Tinney et al., 2024). A key advantage of the technique is in its ability to identify fibre tract-specific changes, overcoming some of the key limitations to DTI and voxel-aggregate scalars (Mito et al., 2018; Raffelt et al., 2017).

Despite the technique’s growing popularity, FBA studies have thus far been largely limited to single-site designs. A key barrier for multi-site diffusion MRI studies more broadly is that quantitative measures are highly susceptible to differences between scanners and protocols, which can pose a major issue when pooling data across sites (Pinto et al., 2020). FBA poses additional challenges for data harmonisation, as recommended pipelines necessitate the construction of study-specific templates, and CSD modelling approaches may differ based on acquisition (e.g., multi-shell vs single-shell) (Dhollander & Connelly, 2016; Jeurissen et al., 2014). Although the extent of scanner or protocol related variability has been well documented for other DWI-based measures (Besseling et al., 2012; Cai et al., 2021; Fan et al., 2021; Grech-Sollars et al., 2015; Koller et al., 2021; Luque Laguna et al., 2020; Schilling et al., 2021; Vollmar et al., 2010), the extent to which site or scanner differences impact fixel-based measures is unclear.

The goal of this study was to assess the reproducibility and reliability of fixel-based measures using participants scanned across multiple sites (a ‘travelling heads’ study). Here, we present the TRAMFIX (TRavelling Across Melbourne for FIXel-based analysis) cohort, in which ten healthy adults were scanned across four 3 T MRI scanners located at three different sites with an identical DWI protocol. We computed fixel-based metrics using two pipelines that could be used when combining data across sites (site-specific versus pooled processing). Reproducibility and reliability of these measures across sites were assessed at three different levels of analysis: (i) averaged across the whole-brain; (ii) averaged across white matter tracts; and (iii) at the individual fixel-level. While the terms reproducibility and reliability are often used interchangeably, they represent different properties (Shoukri et al., 2008), and were assessed using within-subject coefficient of variation (CV_ws_) and intraclass correlation coefficient (ICC), respectively. For comparison, we also examined the reproducibility and reliability of tensor-based metrics at these equivalent three levels (whole-brain, tract-averaged, and voxel-level).

## 2. Methods

### 2.1 Participants

Ten neurologically healthy adult volunteers (7 female, age range: 19-30 years) were recruited into the study via internal email advertisement and notice boards at participating institutions (University of Melbourne, the Florey Institute of Neuroscience and Mental Health, and Monash University). All volunteers were screened for MRI safety and enrolled into the study if they had no contraindications to MRI, and were provided monetary compensation for travel and participation. Participants were included in the analysis if all MRI scans were acquired within a 6-month period. All scans were completed between September to December 2024, with an average interval between first and last scan of 17.8 days (range: 2 to 64 days). Informed written consent was obtained from all participants for being included in the study, and ethical approval was granted by the University of Melbourne’s Human Research Ethics Committee (HREC #28396).

### 2.2 MRI scanner characteristics

Brain images were acquired on four 3T MRI scanners at three different sites across Melbourne, Australia (see Table 1). These included a 3T Magnetom Skyra, Vida, and two Prisma Fits (Siemens Healthcare, Erlangen, Germany). All systems were equipped with a 32-channel head coil.

**Table 1:** Site and scanner information.

|  | *Scanner 1* | *Scanner 2* | *Scanner 3* | *Scanner 4* |
| --- | --- | --- | --- | --- |
| *Site* | Florey | Florey | MBI | RCH |
| *Vendor* | Siemens | Siemens | Siemens | Siemens |
| *Model* | Prisma Fit | Vida | Skyra | Prisma Fit |
| *Maximal gradient amplitude* | 80 mT/m | 60 mT/m | 45 mT/m | 80 mT/m |
| *Software version* | VE11C | XA50 | VE11C | XA30 |
| *Head coil* | 32 ch | 32 ch | 32 ch | 32 ch |

Florey: The Florey Institute of Neuroscience and Mental Health, Melbourne, Australia; MBI: Monash Biomedical Imaging, Monash University, Melbourne, Australia; RCH: Royal Children’s Hospital, Melbourne, Australia.

### 2.3 MRI data acquisition

Each MRI session lasted 1 hour, with approximately 40-50 mins of scan time at each site. Diffusion-weighted images and structural images were acquired at each site, with all sites acquiring one consistent harmonisation protocol for DWI, along with additional site-specific protocols not used in the present study.

The main harmonisation protocol was a multi-shell DWI acquired with a spin-echo EPI sequence (TE/TR = 113/3345 ms, voxel size = 2 mm^3^, matrix size:116 x 116 x 76, multi-band factor 4, no in-plane parallel acceleration). The complete diffusion gradient table consisted of: *b*=0 s/mm^2^ (10 volumes), *b*=300 s/mm^2^, (13 directions), *b*=1000 s/mm^2^ (24 directions), *b*=2000 s/mm^2^ (42 directions), and *b*=3000 s/mm^2^ (60 directions). The diffusion encoding scheme was optimally split, with half the volumes acquired in the anterior-posterior (AP) phase-encoding direction, and half the volumes acquired in the posterior-anterior (PA) direction (designed to achieve optimal sampling across the AP and PA directions using a similar implementation originally developed for the developing Human Connectome Project (dHCP) (Tournier et al., 2020)). Acquisition time for this protocol was ∼9 minutes.

### 2.4 DWI data processing

Here, we discuss the processing steps for the common DWI harmonisation protocol.

All DWI data were preprocessed according to the recommended MRtrix3 pipeline for multi-tissue fixel-based analysis. This included MP-PCA denoising data (Veraart et al., 2016), Gibbs ringing removal (Kellner et al., 2016), EPI susceptibility and motion correction using FSL topup and eddy (Andersson et al., 2003; Andersson & Sotiropoulos, 2016) with eddy QC (Bastiani et al., 2019), bias field correction using ANTs N4ITK (Tustison et al., 2010), and then upsampling all data to a voxel resolution of 1.25 mm^3^. For susceptibility field estimation, only leading *b*=0 volumes were extracted and included, as interleaved *b*=0 volumes were observed to contain residual eddy current distortions from prior volumes.

To mimic two potential strategies for data analysis when combining DWI data across sites for fixel-based analysis, we then processed the cohort using two distinct pipelines (see Figure 1). The first pipeline (Pipeline 1: “site-specific processing”) processed DWI data for each site independently, only pooling data across sites for final analysis. The second pipeline (Pipeline 2: “pooled processing”) instead pooled DWI data across all sites for processing and analysis.

**Figure 1:**
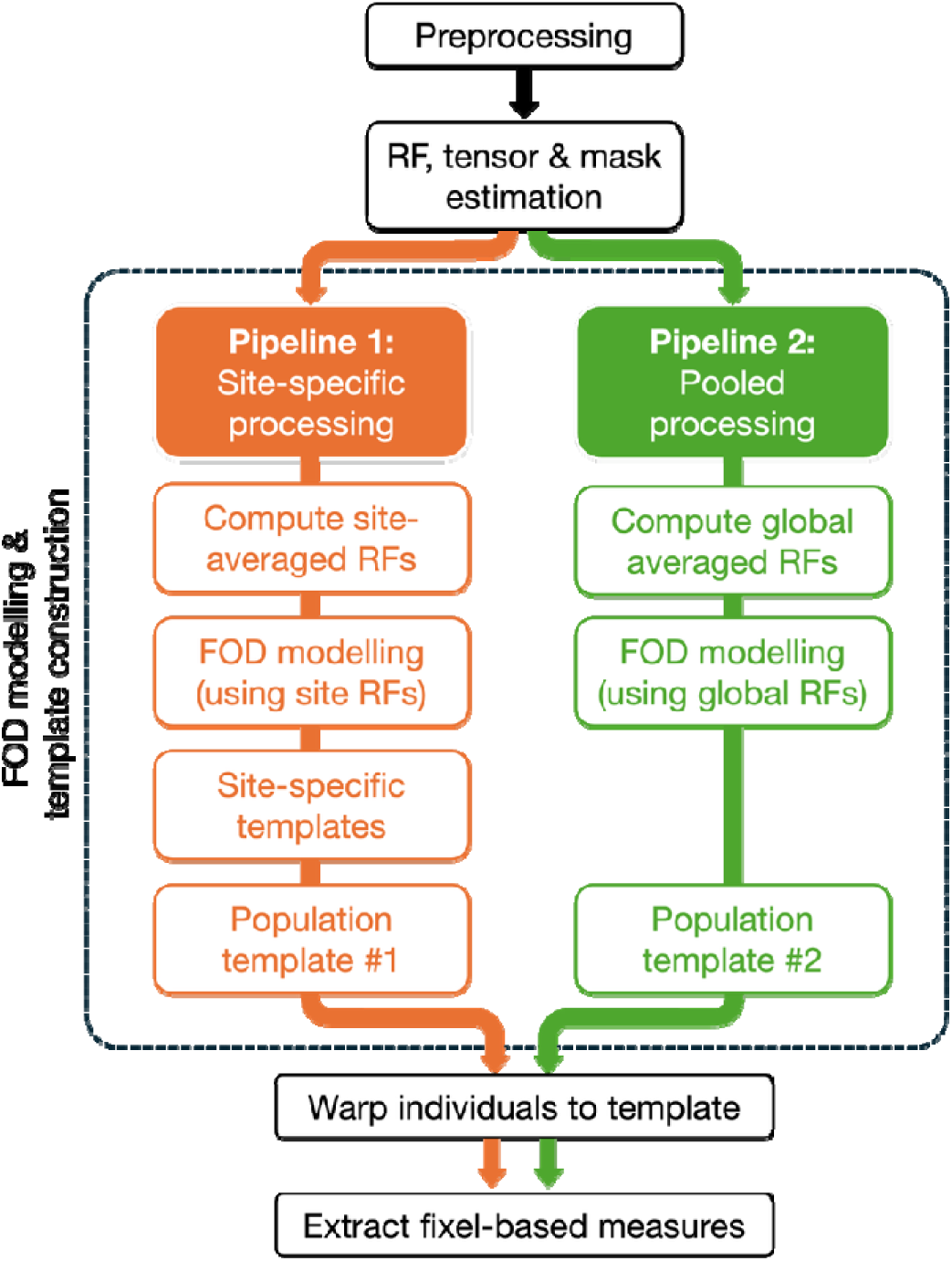
Processing pipelines for extracting fixel-based metrics. Here, we used two alternative approaches that might be taken when combining multi-site data. All data preprocessing, tensor fitting, brain mask estimation, and subject-specific response function estimation were performed in common (once per scanning session). Pipeline 1 describes the scenario where FOD modelling might be performed at a site-specific level (i.e., using site-averaged response functions (RFs) and site-specific templates). Pipeline 2 describes the scenario where data are pooled from different sites prior to FOD modelling (i.e., using globally averaged RFs and a global population template).

In both cases, after DWI data preprocessing, response functions were estimated for white matter, grey matter and CSF (Dhollander et al., 2019) for each participant. Fibre orientation distribution (FOD) functions were then estimated using multi-shell multi-tissue CSD (MSMT-CSD) (Jeurissen et al., 2014). For the first pipeline, site-specific response functions (RFs) were computed (average response functions across all participants at a given site) and used for FOD estimation, while in the second pipeline, response functions averaged across all participants and scanning sessions (i.e., incorporating data from all individuals across all sites) were used for FOD modelling. For each pipeline, a study-specific FOD population template was built using an iterative registration and averaging approach (Raffelt et al., 2011). For Pipeline 1, this was done by first building site-specific templates, then a common population template space using site-specific templates as input; for Pipeline 2, FOD images from all scanning sessions were used in a single template construction step. Each individual FOD image was then registered to the template using FOD-guided non-linear registration (Raffelt et al., 2012).

Measures of fibre density (FD), fibre cross-section (FC), and fibre-density and cross-section (FDC) were then computed according to the fixel-based analysis pipeline (Raffelt et al., 2017). Whole-brain tractography was performed on the population template image, by first generating 20 million streamlines, which were subsequently filtered to 2 million streamlines by performing SIFT (Smith et al., 2013). Smoothing was applied to fixel-based measures using the fixel-fixel connectivity matrix computed from this tractogram (Raffelt et al., 2015).

To compare the reproducibility of fixel-based measures with more conventional tensor-based metrics, we additionally computed measures of fractional anisotropy (FA) and mean diffusivity (MD) from the diffusion tensor. Here, the *b*=1000 s/mm^2^ shell was extracted from the multi-shell harmonisation dataset, and the tensor fit performed using an iterated weighted least-squares approach (Veraart et al., 2013). FA and MD were estimated for each image, and then transformed to population template space (using non-linear transformations computed through registration of FOD images). In order to compare reproducibility of FA and MD to the fixel-based measures, voxel masks were computed as the set of all voxels containing at least one fixel, and tensor measures averaged within those voxel masks. Given that these masks are more extensive than what is typically included within DTI-based analyses, we additionally computed the mean values of these tensor measures within a DTI-specific white matter mask, including only template voxels with a mean FA > 0.2.

We additionally investigated the reproducibility and reliability of fixel-based measures when computed from single-shell data. We extracted *b* = 1000, 2000, and 3000 s/mm^2^ shells from the full multi-shell dataset after preprocessing. Response functions were estimated for each *b*-shell, followed by FOD estimation using single-shell 3-tissue CSD (SS3T-CSD; using MRtrix3Tissue) (Dhollander & Connelly, 2016). As we used the same population template image for these analyses and the fibre cross-section (FC) metric is a measure of deformation to template space, we examined only the reproducibility of the fibre density (FD) measure.

All processing steps (unless otherwise specified) were performed using MRtrix3 (version 3.0.4) (Tournier et al., 2019).

### 2.5 Tract segmentation

Tracts of interest were delineated by performing TractSeg (Wasserthal et al., 2018) on the population template FOD peaks image. Mean FD, FC, and FDC were computed for each tract, including only fixels with correspondence to the local tract direction. Mean voxel-based FA and MD were also computed for each tract, averaging across those voxels containing a fixel ascribed to the tract.

### 2.6 Statistical analysis

To assess the reproducibility and reliability of FBA and tensor-based metrics across the different sites, we computed the coefficients of variation (CV) and intraclass correlation coefficient (ICC).

CV is commonly used to assess the reproducibility of measures, and is broadly defined as the standard deviation of the group (σ) divided by the mean of the group (μ) (often expressed as a percentage). When assessing CV across sites with the same subjects, the within-subject variability (CV_ws_) is considered a more appropriate measure for reproducibility than pooled CV (Shoukri et al., 2008).

CV_ws_ was calculated following recommendations from the quantitative imaging biomarkers alliance (QIBA) for DWI biomarkers (Shukla-Dave et al., 2018). Here, squared CV_ws_ was computed for each subject (dividing variance by squared mean), and average squared CV_ws_ across subjects computed. CV_ws_ was therefore expressed as:

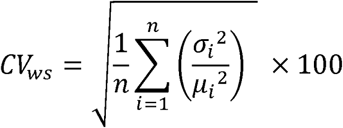

While we focus here on within-subject variability to assess reproducibility, we additionally computed between-subject CV (CV_bs_), which captures biological variability between subjects. CV_bs_ was computed by first taking the within-subject means (*μ_i_*), then computing

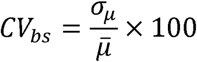

Intraclass correlation coefficient was computed using ICC(3,1), representing two-way mixed effects, consistency, single rater/measurement (Koo & Li,.2016), calculated as: Intraclass correlation coefficient was computed using ICC(3,1), representing two-way mixed

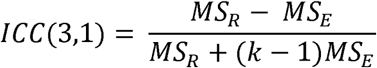

MS_R_ is the mean square for rows (i.e., subjects), while MS_E_ is the mean square for error. While CV_ws_ captures the *reproducibility* of metrics, ICC was used to assess *reliability* of these measures (the overall consistency of measures across individuals) (Luque Laguna et al., 2020; Shoukri et al., 2008). Lower CV_ws_ values reflect better reproducibility, while higher ICC values reflect higher reliability. Generally, ICC values less than 0.5 indicate poor reliability, values between 0.5 and 0.75 reflect moderate reliability, values between 0.75 and 0.9 indicate good reliability, and values greater than 0.9 reflect excellent reliability (Koo & Li, 2016).

CV_ws_ and ICC were assessed for each of the fixel-based measures (FD, FC, and FDC) at: (i) the whole brain level, where measures were averaged across all white matter fixels; (ii) at the tract level, where measures were averaged across tracts of interest; and (iii) at the fixel level, where reproducibility and reliability were assessed at each fixel in template space. These metrics were also computed for tensor-based scalars (FA and MD) at the same three levels.

To compute estimates of sample size requirements for tract-level analyses, we performed power calculations (at α = 0.05, power = 0.8) at different expected observable effect sizes (Cohen’s *d*), using a correction formula implemented in adjusting for group-level correlations (Bliese, 1998), where:

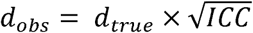

Here, ICC was used as a proxy for reliability, and the computed ICC values from the tract-level results were used. Three specific tracts (corticospinal tract (CST), arcuate fasciculus (AF), and fornix (FX)) were selected *a priori* for sample size calculation to capture different types of tracts. This was based on inclusion in previous repeatability work (Koller et al., 2021). This selection includes commonly studied tracts due to their clinical relevance (CST and AF), along with a tract that is more technically challenging to reconstruct consistently (FX).

## 3. Results

### 3.1 Reproducibility and reliability of fixel-based measures

#### 3.1.1 Comparing whole-brain averaged measures

Table 2 shows the reproducibility (indexed by CV_ws_) and reliability (indexed by ICC) of fixel-based measures (as well as DTI-based measures; see Section 3.2) when averaged across the whole-brain white matter. Across both pipelines, these whole-brain averaged fixel-based measures demonstrated significant evidence of reliability (ICC p-value < 0.0001), with FD, FC, and FDC showing excellent reliability (ICC(3,1) > 0.9 across both pipelines). Variability within subject (CV_ws_) was lower than between subject (CV_bs_) for all measures, ranging between 0.51% to 1.57% for within-subject, and 2.26% to 8.27% for between-subject variability. Reproducibility and reliability metrics were highly comparable between the two pipelines (Table 2).

**Table 2:**
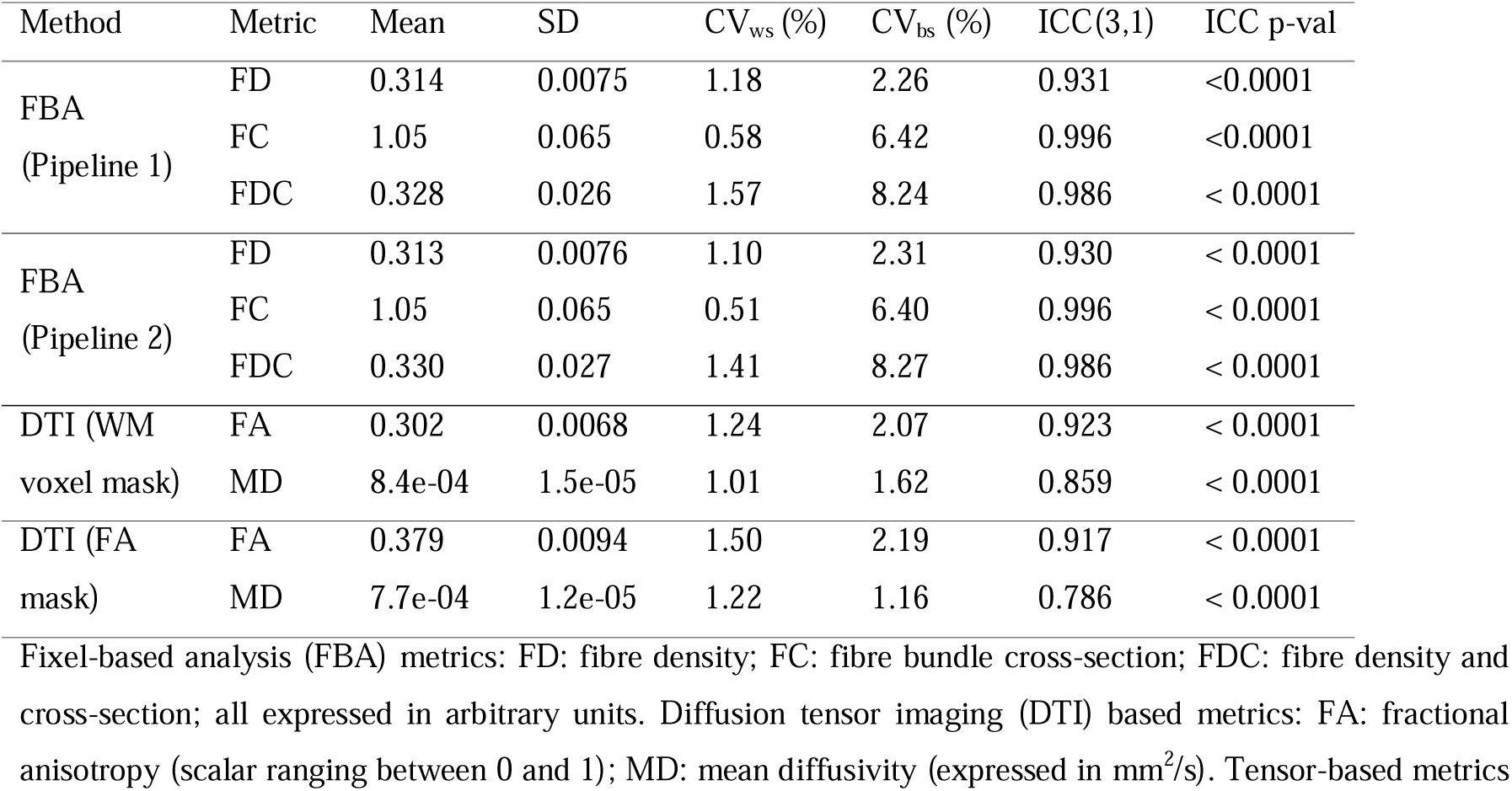
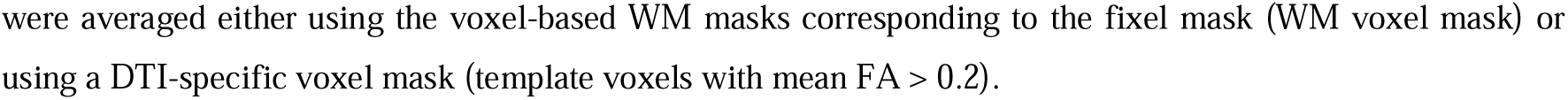
Reproducibility and reliability of fixel-based and DTI-based measures across the whole brain white matter.

| Method | Metric | Mean | SD | $CV_{ws}$ (%) | $CV_{bs}$ (%) | ICC(3,1) | ICC p-val |
| --- | --- | --- | --- | --- | --- | --- | --- |
| FBA<br>(Pipeline 1) | FD | 0.314 | 0.0075 | 1.18 | 2.26 | 0.931 | $<0.0001$ |
| | FC | 1.05 | 0.065 | 0.58 | 6.42 | 0.996 | $<0.0001$ |
| | FDC | 0.328 | 0.026 | 1.57 | 8.24 | 0.986 | $<0.0001$ |
| FBA<br>(Pipeline 2) | FD | 0.313 | 0.0076 | 1.10 | 2.31 | 0.930 | $<0.0001$ |
| | FC | 1.05 | 0.065 | 0.51 | 6.40 | 0.996 | $<0.0001$ |
| | FDC | 0.330 | 0.027 | 1.41 | 8.27 | 0.986 | $<0.0001$ |
| DTI (WM<br>voxel mask) | FA | 0.302 | 0.0068 | 1.24 | 2.07 | 0.923 | $<0.0001$ |
| | MD | 8.4e-04 | 1.5e-05 | 1.01 | 1.62 | 0.859 | $<0.0001$ |
| DTI (FA<br>mask) | FA | 0.379 | 0.0094 | 1.50 | 2.19 | 0.917 | $<0.0001$ |
| | MD | 7.7e-04 | 1.2e-05 | 1.22 | 1.16 | 0.786 | $<0.0001$ |

Fixel-based analysis (FBA) metrics: FD: fibre density; FC: fibre bundle cross-section; FDC: fibre density and cross-section; all expressed in arbitrary units. Diffusion tensor imaging (DTI) based metrics: FA: fractional anisotropy (scalar ranging between 0 and 1); MD: mean diffusivity (expressed in $mm^2/s$ ). Tensor-based metrics
were averaged either using the voxel-based WM masks corresponding to the fixel mask (WM voxel mask) or using a DTI-specific voxel mask (template voxels with mean FA > 0.2).

Figure 2 shows boxplots of the mean fixel-based measures across the four sites.

**Figure 2:**
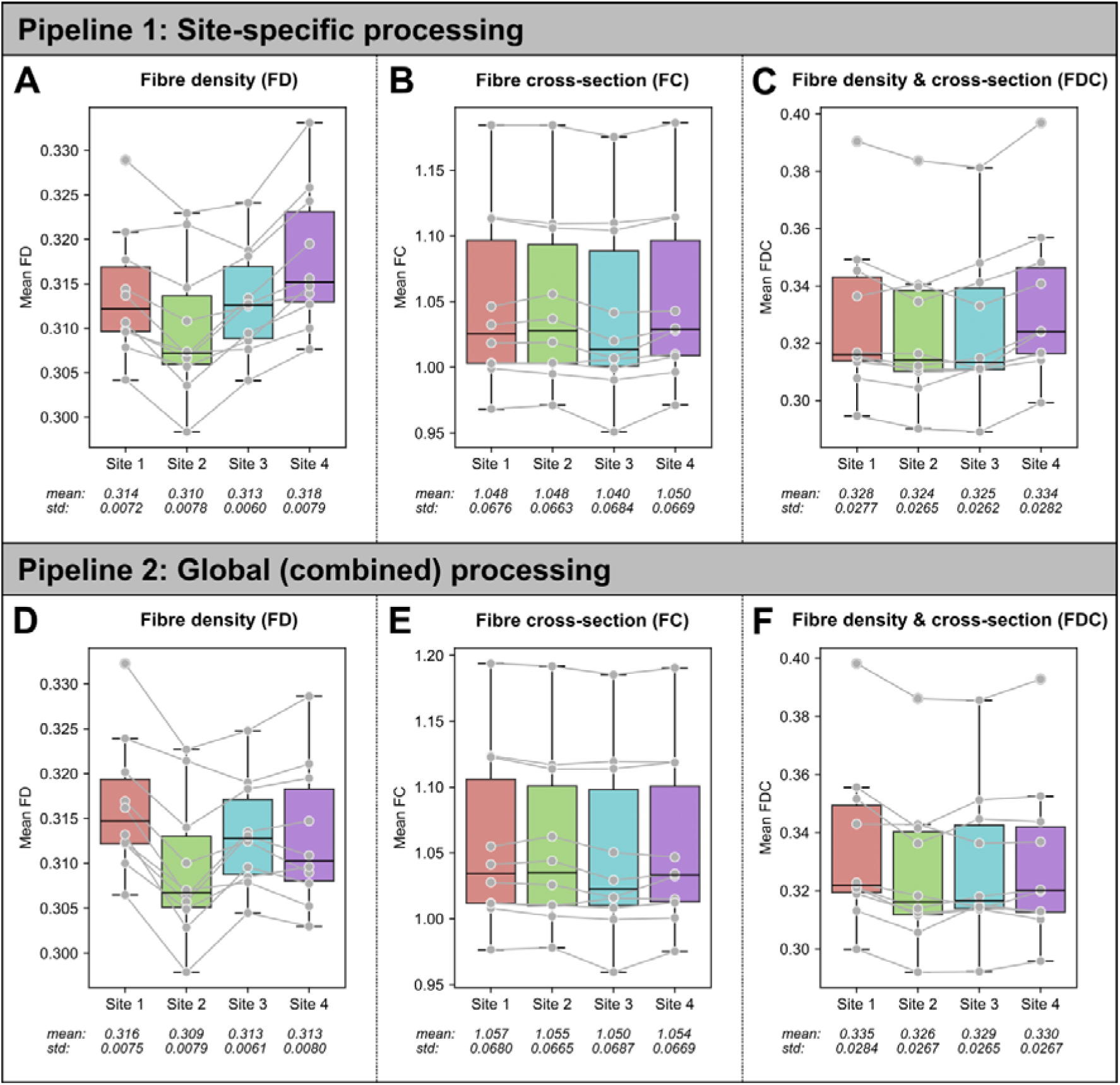
Reproducibility of mean fixel-based measures across the whole-brain. Boxplots show whole-brain fixel-wise measures: (A and D) fibre density (FD); (B and E) fibre cross-section (FC); (C and F) fibre density and cross-section (FDC). The top row shows results for Pipeline 1 (site-specific processing), and bottom row shows results for Pipeline 2 (pooled processing). Points (in grey) represent whole-brain fixel-wise averages for each participant, with grey lines connecting points from the same individual across the four sites. Site-specific mean values and standard deviations (std) are shown for each measure.

#### 3.1.2 Comparing tract-level measures

When comparing tract-averaged fixel-based measures, reproducibility and reliability remained high across tracts. Reproducibility was high for the FD measure for both Pipeline 1 (CV_ws_ ranged between 0.74 – 3.63%) and Pipeline 2 (CV_ws_ ranged between 0.78 – 4.01%). Similarly, reproducibility was high for the FC metric (Pipeline 1 CV_ws_: 0.54 – 1.87%; Pipeline 2 CV_ws_: 0.60 – 1.76%) and for the FDC metric (Pipeline 1 CV_ws_: 1.23 – 5.03%; Pipeline 2 CV_ws_: 1.22 – 5.66%).

Of the fixel-based measures, reliability was most variable for the FD measure (Pipeline 1: ICC ranged between 0.652 – 0.980; Pipeline 2: ICC ranged between 0.633 – 0.980). Reliability was high across both pipelines for the FC measure (Pipeline 1 ICC: 0.966-0.988; Pipeline 2 ICC: 0.967-0.998), and the FDC measure (Pipeline 1 ICC: 0.916-0.995; Pipeline 2 ICC: 0.911-0.994).

Figure 3 shows the ICC values for the FD measure for each white matter tract with Pipeline 1. We focus here on the FD metric, as it was the most affected by site of the fixel-based metrics. Despite the FD measure having the largest variability in reliability, all but two tracts had an ICC > 0.8 (with 65 of 72 tracts showing excellent reliability with ICC > 0.9). Supplementary Figure S1 shows the ICC per tract for the FD measure with Pipeline 2 (same as Figure 3, but with Pipeline 2).

**Figure 3:**
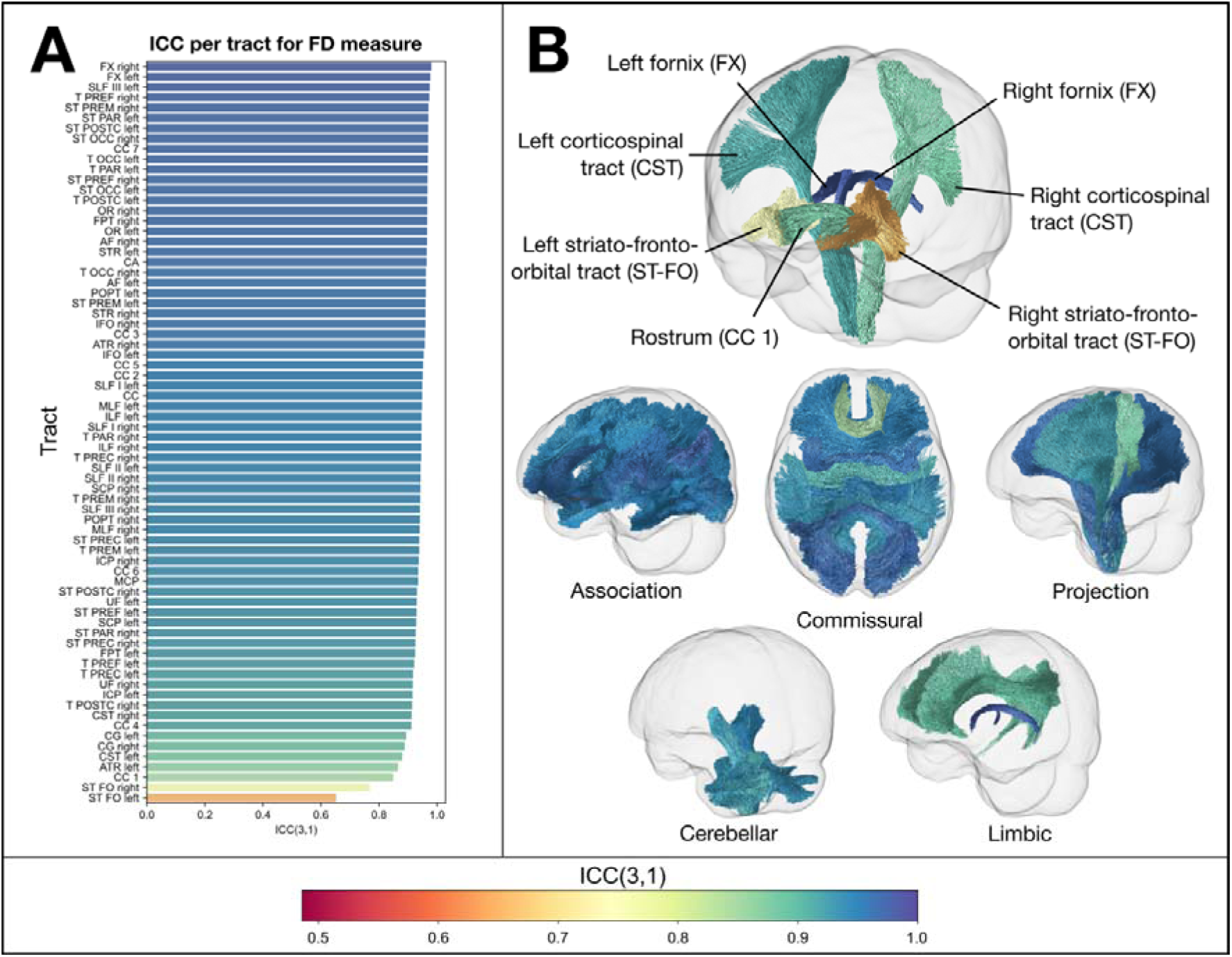
Intraclass correlation coefficient (ICC(3,1)) for the FD measure per tract using Pipeline 1. (A): Bar plot showing ICC for each tract (ordered from highest to lowest). Most tracts showed a very high ICC (> 0.9), with the bilateral fornices (FX) showing the highest of all tracts, and bilateral striato-fronto-orbital tracts (STFO) showing the lowest ICC values. (B): Tract trajectories coloured by their ICC. The top image shows a number of key tracts, showing high (FX), moderate (corticospinal tracts; CST), and lower ICC (striato-fronto-orbital; ST-FO and rostrum of corpus callosum; CC 1). The bottom images show different ICC values for different tract classifications. See Wasserthal et al. (2018) for full tract definitions.

Supplementary Tables S1-S3 report the reproducibility and reliability of fixel-based measures for each white matter tract with Pipeline 1 (site-specific processing).

#### 3.1.3 Comparing fixel-level measures

Figure 4 shows the ICC for each individual fixel in template space (Pipeline 1), for each of the fixel-based measures. In general, reliability was high, with mean ICC across all fixels being 0.83 for FD, 0.95 for FC, and 0.86 for FDC. Figure 5 shows a thresholded image to highlight regions with poor reliability for both FBA and tensor-based metrics (for which results are reported in 3.2). Fixels with lower reliability (ICC < 0.5) tended to be in cerebellar and subcortical grey matter regions, rather than in core white matter tracts. ICC was similarly high at the fixel-level when computing FODs using globally averaged response functions (Pipeline 2), with very comparable results to Pipeline 1 (see Table 3 for summary statistics on ICC across different pipelines and metrics).

**Figure 4:**
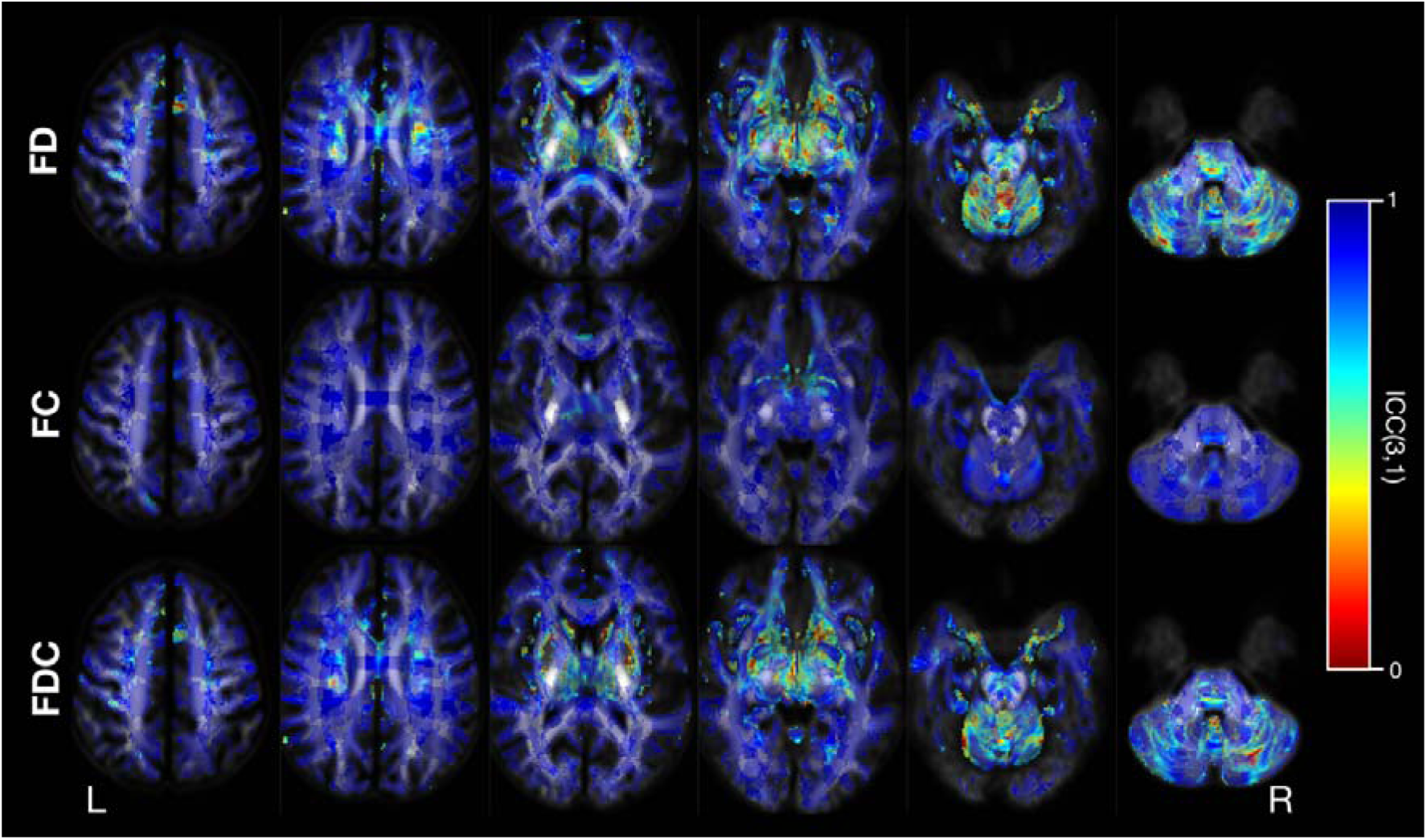
Reliability of FBA metrics at the fixel-level (Pipeline 1). Images show axial slices at 15 mm increments, with white matter fixels overlaid on the FOD template. Fixels are coloured by their ICC value. Top row shows ICC for the FD measure; middle row shows ICC for the FC measure; bottom row shows ICC for the FDC measures.

**Figure 5:**
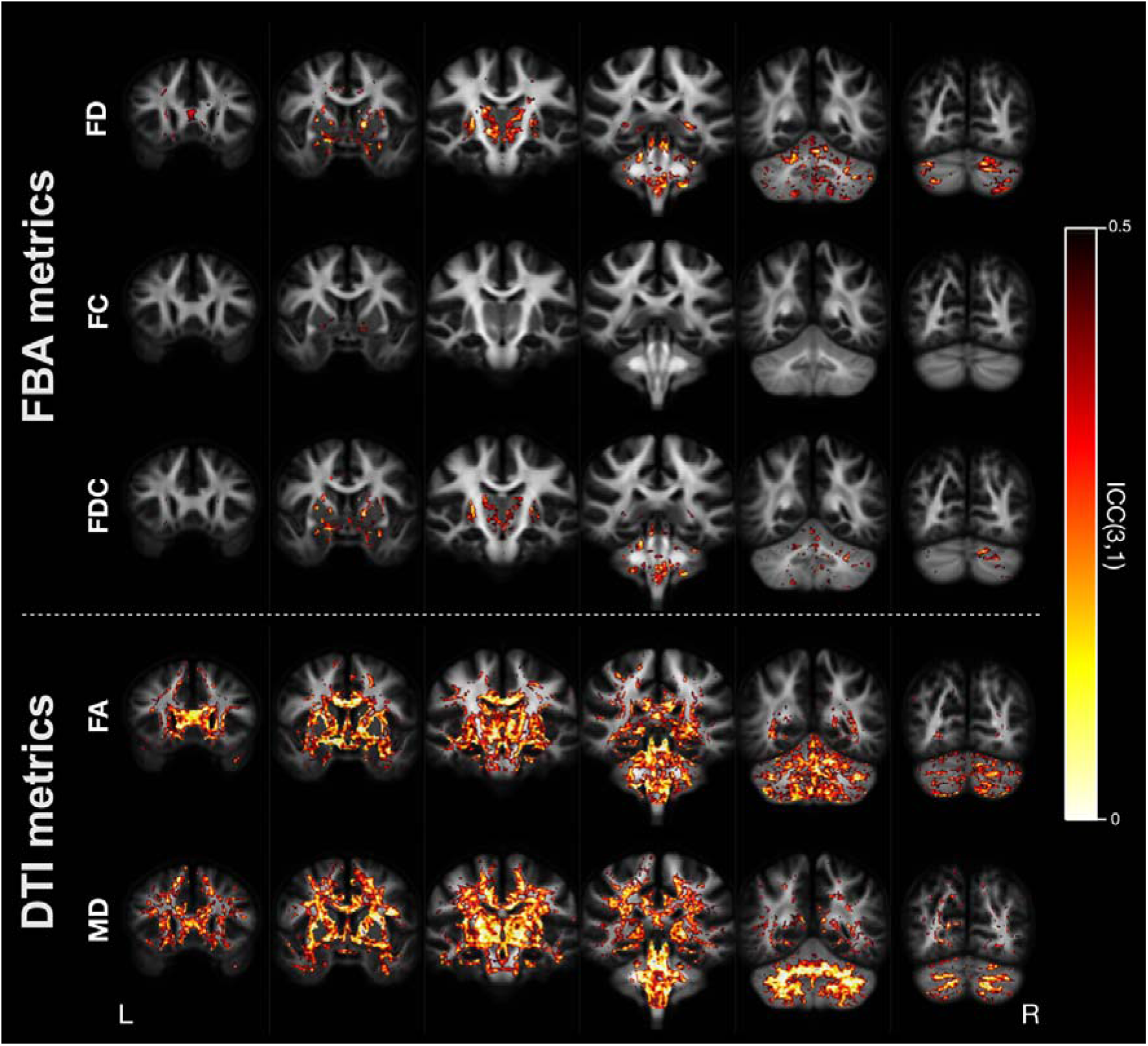
Thresholded ICC image showing regions of lower reliability. ICC is shown for fixel-based analysis (FBA) metrics (top) and tensor-based metrics (bottom). In both cases, the ICC images are thresholded to only show voxels with poor reliability (ICC < 0.5), with bright areas corresponding to lower ICC values and displayed on coronal slices at 18mm increments. For FBA metrics, each voxel is coloured by the minimum value of ICC across the fixels within that voxel (to enable easier visual comparison with tensor-based metrics).

**Table 3:** Summary statistics for ICC at the fixel- or voxel-level.

| Method | Metric | Count | ICC |  |  |  |  |
| --- | --- | --- | --- | --- | --- | --- | --- |
|  |  |  | Mean | Median | SD | Min | Max |
| FBA<br>(pipeline 1) | FD | 627896 | 0.831 | 0.904 | 0.182 | -0.2875 | 0.99996 |
|  | FC | 627896 | 0.954 | 0.970 | 0.049 | 0.0135 | 0.99801 |
|  | FDC | 627896 | 0.863 | 0.923 | 0.154 | -0.2920 | 0.99952 |
| FBA<br>(pipeline 2) | FD | 625296 | 0.832 | 0.905 | 0.183 | -0.2861 | 0.99969 |
|  | FC | 625296 | 0.955 | 0.971 | 0.049 | 0.1624 | 0.99842 |
|  | FDC | 625296 | 0.863 | 0.925 | 0.156 | -0.254 | 0.99969 |
| DTI (WM<br>voxel mask) | FA | 431463 | 0.596 | 0.646 | 0.252 | -0.2999 | 0.99493 |
|  | MD | 431463 | 0.551 | 0.589 | 0.262 | -0.2931 | 0.99212 |
| DTI (FA<br>mask) | FA | 284920 | 0.590 | 0.637 | 0.253 | -0.2999 | 0.99446 |
|  | MD | 284920 | 0.484 | 0.504 | 0.258 | -0.2931 | 0.99212 |

Table 4 shows the reproducibility (within-subject variability; CV_ws_) of fixel-based measures, summarised across all fixels within the white matter mask (density distribution plots shown in Supplementary Figure S3). Overall, mean CV_ws_ was lowest for the FC measure (mean CV_ws_ < 3%), while substantially higher for the FD and FDC measures. Higher average CV_ws_ across all fixels for these two measures were likely driven by very large variability in a small number of fixels, with the mean CV_ws_ around 15%, and median CV_ws_ around 10% across all fixels. Figure 6 shows CV_ws_ for each of the metrics on a single axial slice. Core white matter pathways tended to have high reproducibility, while lower reproducibility (higher CV_ws_) was observed in the FD and FDC metrics in fixels closer to the cortical surface, and fixels crossing major white matter regions into subcortical regions (e.g., crossing fibre structures in the anterior limb of the internal capsule).

**Figure 6:**
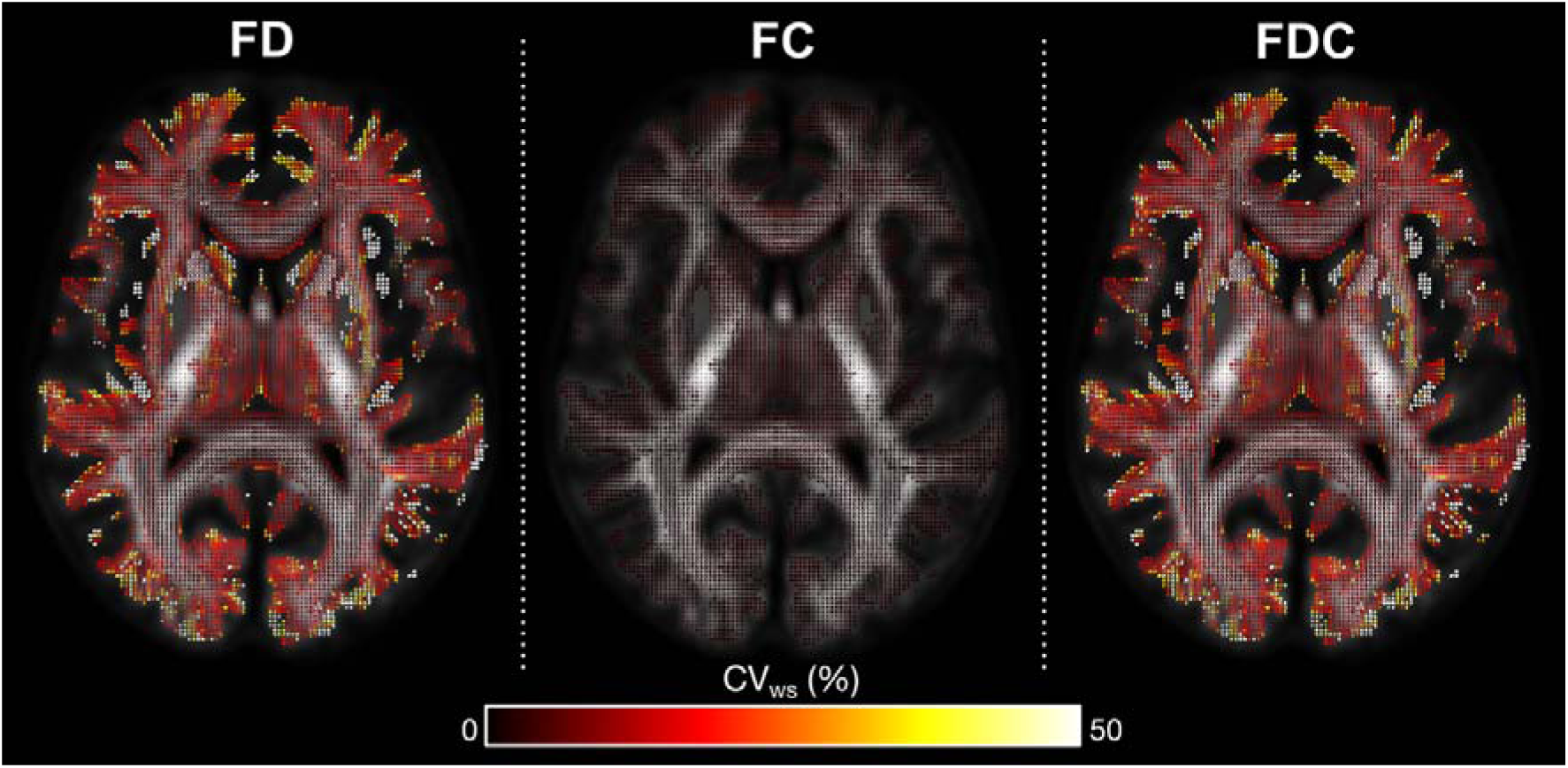
Fixel-level reproducibility. CV_ws_ is shown for each of the fixel-based measures: fibre density (FD), fibre cross-section (FC), and fibre density and cross-section (FDC) on a single axial slice.

**Table 4:** Summary statistics for CV_ws_ at the fixel- or voxel-level.

| Method | Metric | Count | $CV_{ws}$ | | | | |
| --- | --- | --- | --- | --- | --- | --- | --- |
|  |  |  | Mean | Median | SD | Min | Max |
| FBA<br>(Pipeline 1) | FD | 627896 | 15.95 | 10.82 | 18.87 | 0.59 | 200 |
|  | FC | 627896 | 2.27 | 2.11 | 0.79 | 0.74 | 13.11 |
|  | FDC | 627896 | 16.37 | 11.25 | 18.73 | 1.47 | 200 |
| FBA<br>(Pipeline 2) | FD | 625296 | 16.05 | 10.79 | 19.24 | 0.82 | 200 |
|  | FC | 625296 | 2.22 | 2.05 | 0.76 | 0.68 | 7.39 |
|  | FDC | 625296 | 16.45 | 11.22 | 19.11 | 1.53 | 200 |
| DTI (WM<br>voxel mask) | FA | 431463 | 20.43 | 20.62 | 8.84 | 1.75 | 64.37 |
|  | MD | 431463 | 7.04 | 5.85 | 21.93 | 1.43 | 14101 |
| DTI (FA<br>mask) | FA | 284920 | 17.42 | 16.71 | 8.20 | 1.75 | 64.37 |
|  | MD | 284920 | 5.99 | 4.82 | 3.80 | 1.43 | 321.9 |

### 3.2 Reproducibility and reliability of tensor-based measures

In addition to comparing fixel-based measures, we also examined the reliability reproducibility of tensor-based fractional anisotropy (FA) and mean diffusivity (MD). When using a white matter voxel mask equivalent to the fixel mask used for FBA analyses, mean FA and MD showed high reliability and reproducibility, with high ICC (FA ICC = 0.923; MD ICC = 0.859) and low CV_ws_ (FA CV_ws_ = 1.24%, MD CV_ws_ = 1.01%). Reliability and reproducibility were slightly lower when FA and MD were averaged across a constrained WM mask (see Table 2). Boxplots in Supplementary Figure S4 show mean whole-brain FA and MD for participants across sites.

Taking tract-averaged FA and MD using the same TractSeg tract definitions used for FBA metrics, we examined the reproducibility of measures averaged at each white matter tract. Within-subject variability (CV_ws_) for FA ranged from 0.65% to 4.58%, and intraclass correlation coefficient (ICC) from 0.486 to 0.963, with the lowest ICC reported in cerebellar tracts (middle cerebellar peduncle (MCP) and bilateral inferior cerebellar peduncles (ICP)). The CV_ws_ for MD ranged from 0.90% to 1.98%, and ICC between 0.465 and 0.969. Figure 7 shows ICC for each of the TractSeg tracts for the FA measure. In general, reproducibility was lower for the FA measure than for the FD measure, and tracts showing lower/higher reproducibility did not necessarily overlap with the FD measure (see Figure 3 for comparison). Supplementary Tables S4 and S5 show results for each tract for FA and MD, respectively.

**Figure 7:**
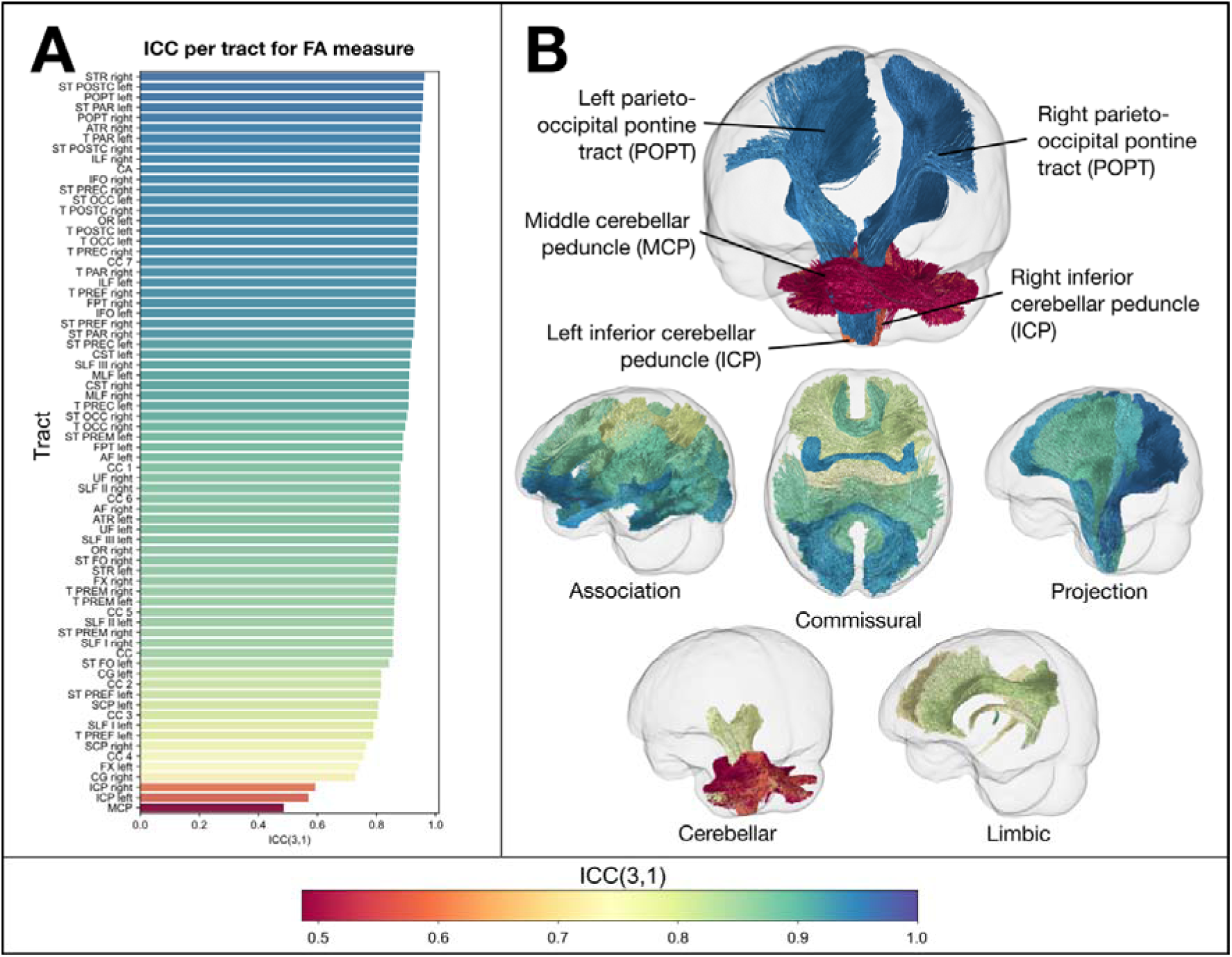
Intraclass correlation coefficient (ICC(3,1)) for the FA measure per tract. (A): Bar plot showing ICC for each tract (ordered from highest to lowest), using same colour scaling as Figure 3. (B): Glass brain images show tracts coloured by their ICC value, for a selection of tracts with the highest and lowest ICC (top), and for different tract classifications (bottom images). Here, reproducibility was lowest for the cerebellar white matter tracts (middle cerebellar peduncle (MCP) and bilateral cerebellar peduncles (ICP)), and highest in the motor system tracts and thalamic radiations.

When examining ICC across white matter voxels independently, a similar pattern was observed to fixel-based measures, with ICC lowest in cerebellar and subcortical white matter regions. However, mean ICC was substantially lower for tensor-based metrics than for fixel-based measures, with mean ICC across all WM voxels at 0.60 for FA and 0.55 for MD (Table 3). Given that the fixel-based white matter mask may be more extensive than white matter masks used for DTI-based analyses, we also computed a DTI-specific white matter mask (constraining to voxels where the FA > 0.2). However, results were similar when using the constrained WM mask, with mean ICC across all WM voxels at 0.59 for FA, and 0.48 for MD. Figure 5 shows voxels with poor reliability for both the fixel-based and tensor-based measures, demonstrating the greater extent of lower reliability voxels in tensor-based metrics, which included some key white matter regions (e.g., corpus callosum, internal capsule).

The reproducibility of tensor-based metrics at the voxel level was generally high within core white matter tracts, with CV_ws_ similar to that of FBA metrics. Supplementary Figure S3 shows distribution of voxel-wise ICC and CV_ws_ for all metrics, while Supplementary Figure S4 shows ICC and CV_ws_ for the FA and MD measures on a single axial slice, alongside whole-brain boxplots.

### 3.3 Reproducibility of fibre density measure at different b-values

The fibre density (FD) measure is known to be dependent on *b*-value, and as such, we also examined whether reproducibility of this measure would be different at different *b*-values (see Supplementary Figure S5). Here, we extracted each *b*-shell from the full multi-shell DWI, and modelled FODs using the single-shell data only. The FD measure was indeed dependent on *b*-value, with higher mean FD across the whole brain at *b*=1000 s/mm^2^ and *b*=2000 s/mm^2^ than at *b*=3000 s/mm^2^. Reproducibility of the FD measure was high across all *b*-shells, though was highest when using the *b*=1000 s/mm^2^ data (ICC of 0.921), followed by the *b*=3000 s/mm^2^ data (ICC of 0.914), and lowest for the *b*=2000 s/mm^2^ shell (ICC of 0.879) (see Supplementary Table S6). Reproducibility of FD was high across all *b*-shells, with all demonstrating within-subject variability (CV_ws_) around 1%.

### 3.4 Sample size estimates for tract-level analyses

Sample size estimates were computed for tract-level analyses, by using the ICC values reported in sections 3.1.2 and 3.2 to estimate observable effect sizes at different expected true effects. Figure 8 shows sample size requirements per group for an independent sample t-test, with a = 0.05 and power = 0.8, for three select tracts (fornix, arcuate fasciculus, and corticospinal tract), for both the fibre density (FD) and fractional anisotropy (FA) metrics across a range of expected true effect sizes (0.5 < d < 1.0). Sample size requirements were similar for fixel-based FD to tensor-based FA, but smaller tracts like the fornix had lower sample size requirements for FD, due to the greater impact of site variability on the FA measure than for the FD measure.

**Figure 8:**
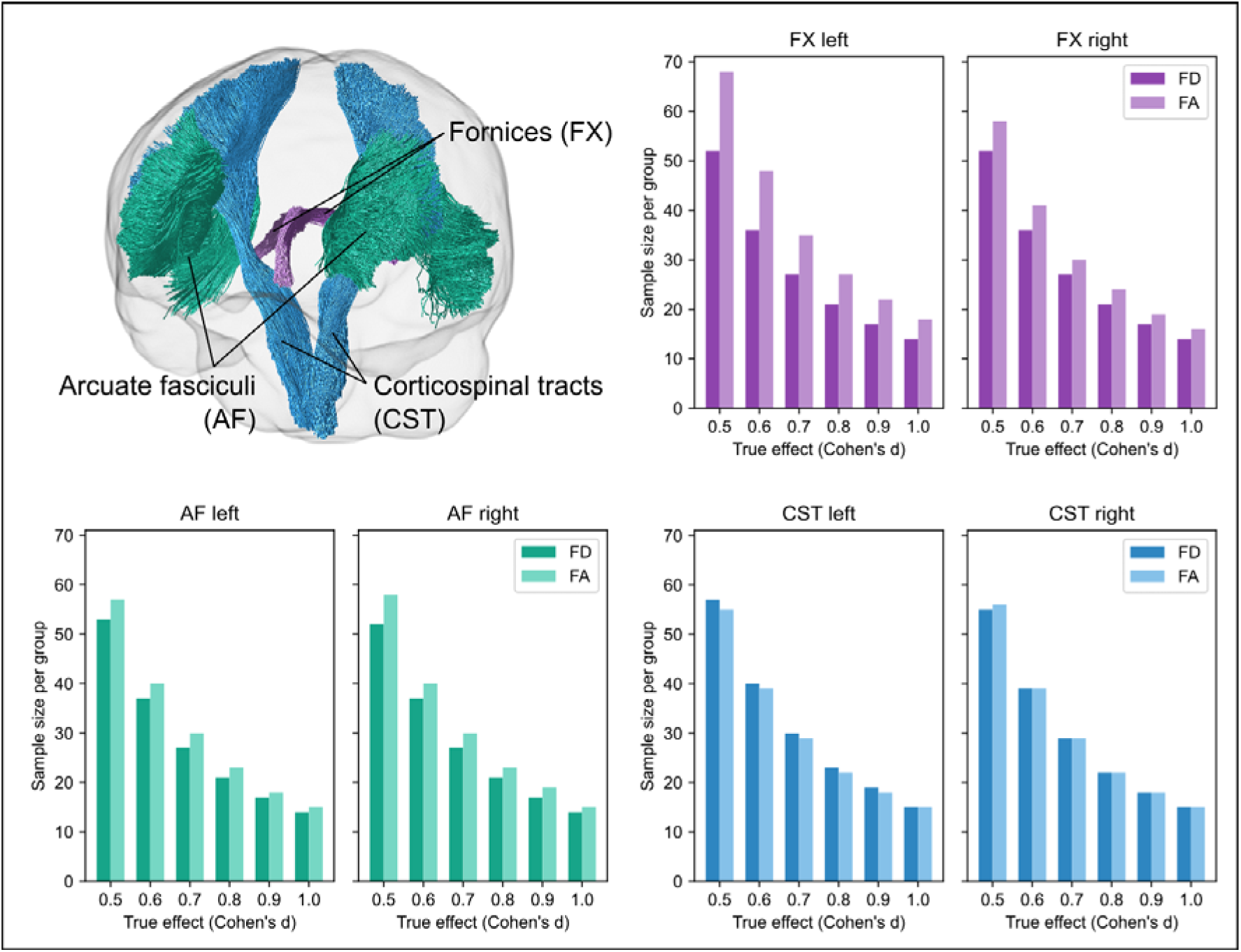
Sample size estimates for tract-level analyses for fibre density (FD) and fractional anisotropy (FA). Glass brain in top left shows the three tracts selected for sample size estimates (fornix (FX); arcuate fasciculus (AF); corticospinal tract (CST)). Bar plots show sample size requirements for each of these tracts, using the ICC values that were computed from the tract-level reproducibility analyses. Given the lower reproducibility of the FA measure for the fornix, sample size requirements to detect significant effects in this tract were higher than for the FD measure. For other tracts, sample size estimates were comparable.

Of note, these sample size estimates are based on *independent* tract analyses. More stringent alpha thresholds that might be used when conducting multiple tract analyses would result in higher sample size requirements.

## 4. Discussion

In this work, we present the TRAMFIX dataset, a travelling heads dataset designed to probe the reproducibility and reliability of fixel-based analysis (FBA) metrics derived from DWI data. We demonstrate that whole-brain and tract-averaged FBA measures are highly reproducible and reliable across scanners when using consistent protocols. However, when examining reproducibility and reliability at individual fixels, it was evident that certain white matter regions are more greatly impacted by scanner variability than others, which may be underappreciated at the whole-brain and tract-averaged level. This highlights the importance of careful interpretation in certain brain regions when conducting multi-site studies, and the potential need for appropriate harmonisation approaches. Of note, both reproducibility and reliability of FBA metrics was comparable, if not higher, than DTI-based measures, particularly for small fibre structures like the fornix, and in subcortical and cerebellar white matter. However, we also highlight that the *b*-value dependence of fixel-based measures should be taken into account when pooling data across different protocols. We additionally provide recommendations for sample size estimates when performing tract-level analyses with multi-site datasets, in light of the observed variability. Together, our findings provide novel insight into the reproducibility and reliability of fixel-based measures, which will potentially guide future studies that expand FBA beyond the single-site, and into multi-site designs.

### 4.1 Reproducibility and reliability of FBA measures

There has been growing interest in the repeatability, reproducibility, and reliability of DWI-derived measures, as quantitative MRI measures are known to demonstrate some level of variability even under identical conditions (Jansen et al., 2007; Koller et al., 2021; Vollmar et al., 2010), and to be highly susceptible to differences in scanner hardware and data acquisition (Mirzaalian et al., 2016; Tax et al., 2019). There have now been a number of studies examining the reproducibility of tensor-based metrics (Besseling et al., 2012; Koller et al., 2021; Luque Laguna et al., 2020; Magnotta et al., 2012; Palacios et al., 2017; Vollmar et al., 2010), measures from multicompartment models (Andica et al., 2020; Fan et al., 2021; Veraart et al., 2021), and fibre tractography (Schilling et al., 2021). This work extends our understanding of DWI reproducibility to fixel-based metrics, which are being increasingly adopted in research studies due to their ability to identify fibre tract-specific differences in clinical cohorts. Our findings demonstrate that FBA measures are highly reproducible and reliable across sites when using identical imaging protocols. FBA metrics demonstrated lower within-subject variation (CV_ws_) than between-subject variation (CV_bs_), and high intraclass correlation coefficient (ICC) across the different scanners, with comparable results to previous literature in other DWI metrics. Within our study, we similarly observed comparable reproducibility in FBA metrics as with DTI-based FA and MD, with slightly higher reliability evident for FBA measures particularly at the tract- and voxel-level.

Our findings may be taken to support the expansion of fixel-based analysis studies beyond predominantly single-site into multi-site study designs. Indeed, reliability and reproducibility of fixel-based measures was very high when looking at whole-brain averaged or tract-averaged metrics, with high ICC values and low within-subject variability across most white matter bundles. This was in keeping with other DWI metrics, where inter-scanner variabilities have generally been reported to be less than 10% within select regions or tracts of interest (Cai et al., 2021; Grech-Sollars et al., 2015; Palacios et al., 2017; Vollmar et al., 2010). DTI-based studies have increasingly been expanded to multi-site designs, and the comparable reproducibility of FBA metrics reported here demonstrates promise for studies that may wish to explore fibre-specific white matter differences in cohorts with multi-site data collection. To this end, we provide sample size estimates when studies may wish to probe tract-specific differences in multi-site cohorts. However, we do recommend caution when interpreting findings in brain regions that may be more susceptible to scanner differences.

Importantly, in this study, we additionally examined reliability and reproducibility of fixel-based metrics at the individual fixel-level (as well as DTI-metrics at the voxel-level). Of note, previous studies examining variability of DWI measures across sessions or sites have predominantly focused on quantifying reliability (ICC) or reproducibility (CV_ws_) in whole-brain, tract-, or ROI-averaged metrics. While this is a valuable way to assess and summarise DWI metrics, many clinical research studies aim to conduct group analyses at the more granular voxel-level (or in the case of FBA, at the fixel-level). Studies that have examined variability in DWI metrics at the voxel-level have generally assessed correlation (using Pearson’s R) or absolute deviation between paired tests (e.g., test-retest pairs) (Fan et al., 2021; Koller et al., 2021). Here, we provide ICC and CV_ws_ estimates at each individual fixel or voxel, enabling easy visualisation of brain regions that are affected by scanner differences.

Indeed, our fixel- and voxel-level results indicated clear spatial patterns to reliability, with much lower ICC in some brain regions than was observable at the tract or whole-brain averaged level. The spatial pattern of low ICC within subcortical structures was similar to previous reports that have shown lower ICC in voxel-based measures (Luque Laguna et al., 2020). Areas of low reliability (Figure 5) were predominantly in the subcortex and cerebellum, likely reflecting scanner-related differences in achievable SNR in these regions. However, we also note that for DTI-based measures, lower ICC was observed across many core white matter voxels. Reproducibility tended to be high across most white matter fixels for FBA metrics, and voxels for tensor-based metrics, with greater within-subject variability at cortical boundaries or in minor crossing fixels. Providing estimates of reliability and reproducibility at the individual voxel- or fixel-level may be informative for studies conducting multi-site DWI analyses, particularly when using scanners with different gradient systems.

Of note, we investigated two potential pipelines that might be used when pooling data for fixel-based analyses across sites. Modelling data using constrained spherical deconvolution (CSD) relies on first estimating a response function from the DWI data. As fibre orientation distribution functions are estimated from the derived response functions (by deconvolution of the response function from the measured DWI signal (Tournier et al., 2007), the use of standardised (for example, group-averaged) response functions is recommended when performing fixel-based analysis (Raffelt, Tournier, Rose, et al., 2012; Raffelt et al., 2017). We tested reproducibility of fixel-based measures using both site-specific response functions and FOD estimation, and pooled average response function estimation and FOD estimation. Although we expected combined processing (using pooled average RFs and FOD estimation) would reduce variability and improve reproducibility and reliability across sites, the results were in fact highly similar across the two pipelines. This may suggest that when combining data for FBA across multiple sites, data processing (that is, DWI modelling steps prior to template registration) could reasonably be performed at a site-specific level prior to pooling data across sites for further analysis (coregistration to a common template, and statistical analysis), at least where DWI data have been collected using matched protocols. However, we note that for this particular study, we made significant effort to utilise identical protocols with matched parameters across all sites, while including scanners from the same vendor, resulting in low variability across sites (CV_ws_ < 5% across all white matter tracts). Where there is greater variability across sites, group-based processing may still help to improve reproducibility and reliability of fixel-based measures.

Finally, we note that when assessing reproducibility of diffusion-based measures, coefficient of variation is often used (Cai et al., 2021; Grech-Sollars et al., 2015; Koller et al., 2021; Luque Laguna et al., 2020; Magnotta et al., 2012; Pfefferbaum et al., 2003; Vollmar et al., 2010). Although CV is quite simply defined as the standard deviation of measurements divided by the mean, when used to assess reproducibility across individuals and sites, separating the variability into within-subject and between-subject variability is valuable for differentiating site-related variability (i.e., reproducibility) from between-subject differences (Shoukri et al., 2008). However, even when computing CV_ws_, studies will often used a pooled (grand) mean (Kurokawa et al., 2021; Luque Laguna et al., 2020), which still captures between subject differences. Here, we implement a robust measure of CV_ws_, as recommended by the quantitative imaging biomarkers alliance (QIBA) (Shukla-Dave et al., 2019), and advocate for future studies to similarly quantify reproducibility in this way. We also echo previous recommendations to assess both *reproducibility* (generally indexed with within-subject variability) and *reliability* (indexed with ICC) in such studies, as they reflect separate properties (Shoukri et al., 2008; Luque Laguna et al., 2020).

### 4.2 Limitations and future directions

To our knowledge, this is the first study to probe the reproducibility and reliability of fixel-based analysis (FBA) measures, making use of a travelling heads cohort. Here, we were particularly interested in assessing scanner-related variability across local neuroimaging research centres, and as such, focused on these inter-scanner differences while making use of an identical harmonised DWI protocol across sites (that is, we focused on *reproducibility* rather than *repeatability* (Plesser, 2018; Raunig et al., 2015)). We were unable to assess inter-session (test-retest) repeatability, as well as potential protocol-related variability in fixel-based measures. As has been previously demonstrated (Genc, Tax, et al., 2020), we note the *b*-value dependence of the apparent fibre density (FD) measure, highlighting that protocol-related variability may be much more pronounced than scanner-related differences. We also note that the use of multi-shell versus single-shell data and subsequent FOD modelling could result in further variability in derived metrics. To this end, we hope that the release of the TRAMFIX dataset will enable researchers to additionally characterise protocol-related variability for fixel-based measures.

Another limitation to our work was the inclusion of scanners from the same vendor. Previous work has highlighted that vendor-related variability in diffusion-derived features may be much greater than that from different scanner models of the same vendor (Andica et al., 2020; Schilling et al., 2021). Of note, even with identical protocols and scanner model (Scanner 1 and 4), we observed some variability in derived measures, as has been previously reported for DTI-based metrics. However, we note that future work probing the impact of different vendors on fixel-based measures may also be valuable.

Finally, while we focused in this study on assessing the reproducibility and reliability of fixel-based analysis metrics, to bring FBA studies into multi-site designs, future work evaluating the performance of different harmonisation approaches may be valuable. There are now a range of potential diffusion harmonisation approaches available (Pinto et al., 2020; Tax et al., 2019), including those that harmonise on the diffusion signal (Mirzaalian et al., 2016), and batch harmonisation approaches for derived metrics (Fortin et al., 2017). These may be particularly pertinent when performing whole-brain analyses rather than examining tract-averaged metrics, given the variable reliability across fixels. The extent to which different harmonisation approaches perform on fixel-based measures remains only briefly characterised (Mito et al., 2023). Travelling subject datasets can be valuable in this way, as they enable biological variability to be kept at a minimum, while assessing the performance of different harmonisation approaches. We hope that the TRAMFIX dataset will be a valuable resource to future studies that may wish to assess the performance of harmonisation techniques, and provide this as a resource alongside other travelling or test-retest datasets available in the community (Cai et al., 2021; Koller et al., 2021; Kurokawa et al., 2021; Tax et al., 2019).

### 4.3 Conclusion

Here, we present a travelling heads data for fixel-based metrics – the TRAMFIX dataset (TRavelling Across Melbourne for FIXel-based analysis). We demonstrate high reliability and reproducibility of fixel-based measures across scanners from the same vendor and sites, when using identical protocols, with similar results to that reported from other DWI measures. Future studies that examine the impact of different protocols as well as performance of data harmonisation approaches are likely to be valuable. To this end, we provide the TRAMFIX dataset as a resource for future work. These findings may help to guide large multi-site study designs, and improve the reach of more advanced diffusion approaches like fixel-based analysis.

## Data and Code Availability

The code used to process DWI data, and to perform statistical analyses are available on GitHub (https://github.com/remikamito/TRAMFIX).

## Author Contributions

Conceptualisation: RM, SG

Methodology: RM, RS

Data curation: RM, JH, MK, CK, RM

Investigation: RM, SG, JYMY, RS

Formal analysis: RM

Software: RM, RS

Writing – Original Draft: RM

Writing – Review & Editing: RM, SG, JH, JYMY, JDT, MAD, RS, AZ

Funding acquisition: RM

## Acknowledgments

We are very grateful to the participants for volunteering their time for this study.

The authors acknowledge the facilities and scientific and technical assistance of the National Imaging Facility (NIF), a National Collaborative Research Infrastructure Strategy (NCRIS) capability. The authors also thank the radiographers and facility staff at each of the research sites for their expertise and support: the Florey MRI facility, Monash Biomedical Imaging, and Royal Children’s Hospital. We are also very grateful to Dr Michelle Wu for her assistance in the study.

R.M. is the recipient of an Australian Research Council Discovery Early Career Researcher Award (project number: DE240101035) and University of Melbourne Establishment Grant. S.G. and J.Y-M.Y. acknowledge funding from the Royal Children’s Hospital Foundation (RCHF 2022-1402). S.G. acknowledges support from the Melbourne Children’s Clinician-Scientist Fellowship. J.Y-M.Y. acknowledges support from The Kids’ Cancer Project (TKCP) Col Reynolds Fellowship. JDT is supported by core funding from the Wellcome/EPSRC Centre for Medical Engineering [WT203148/Z/16/Z] and by the National Institute for Health Research (NIHR) Biomedical Research Centre based at Guy’s and St Thomas’ NHS Foundation Trust and King’s College London and/or the NIHR Clinical Research Facility. The views expressed are those of the author(s) and not necessarily those of the NHS, the NIHR or the Department of Health and Social Care. A.Z. is supported by an Australian Research Council Future Fellowship and Rebecca L. Cooper Foundation Fellowship.

