## Supplementary Material for "TRAMFIX: TRavelling Across Melbourne for FIXel-based analysis (a reproducibility study)"

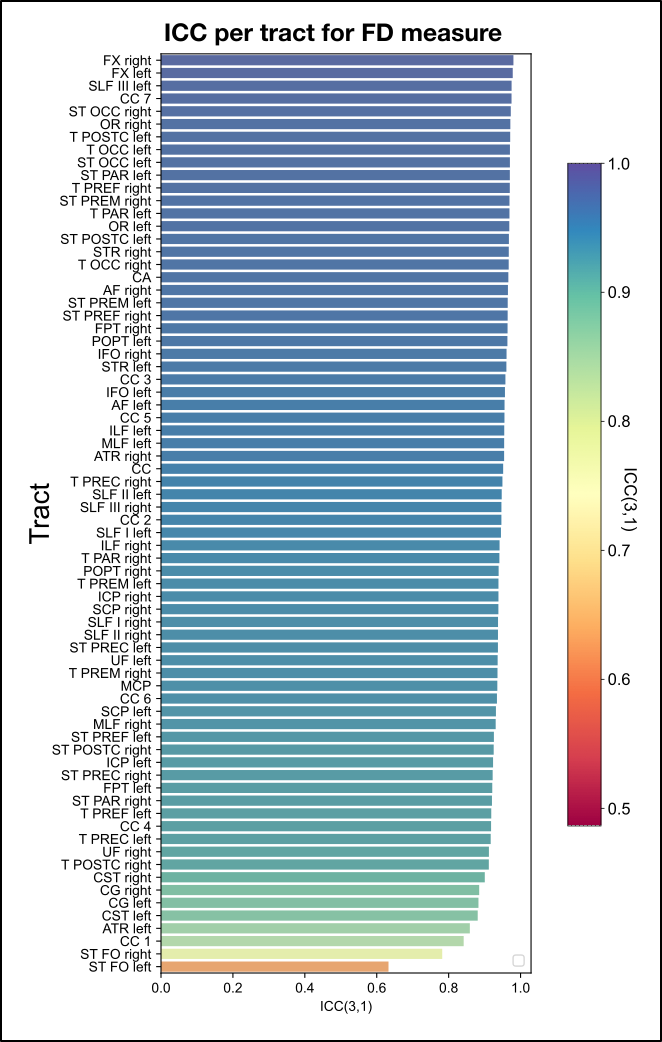

**Supplementary Figure S1: Intraclass correlation coefficient (ICC(3,1)) for the FD measure per tract for Pipeline 2.** Bar plot shows ICC for each tract (ordered from highest ICC to lowest). FD values were computed using Pipeline 2 (group-based processing), which yielded very similar reproducibility results to Pipeline 1 (data shown in Figure 3). As with Pipeline 1, most tracts showed a very high ICC (65/72 tracts with ICC > 0.9). The bilateral fornices (FX) showing the highest of all tracts, and bilateral striato-fronto-orbital tracts (STFO) showing the lowest ICC values. The same colour scale is used here as for Figure 3 and Figure 6 for comparability.

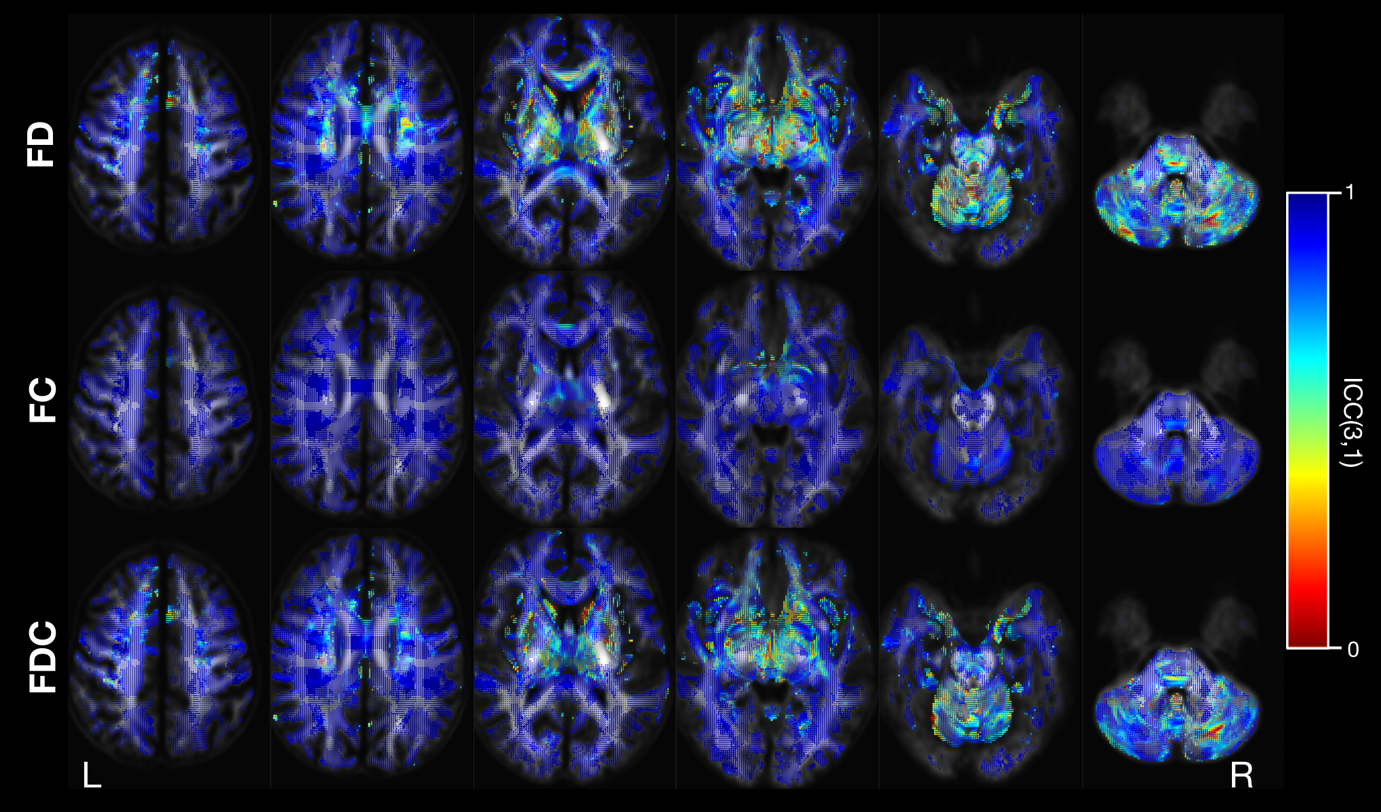

**Supplementary Figure S2: Intraclass correlation coefficient (ICC(3,1) for fixel-based metrics per fixel for Pipeline 2.** Top row shows ICC for the fibre density (FD) metric, middle row shows ICC for the fibre cross-section (FC) metric, and bottom row shows ICC for the fibre density and cross-section (FDC) metric. Results were very similar to Pipeline 1 (Figure 4).

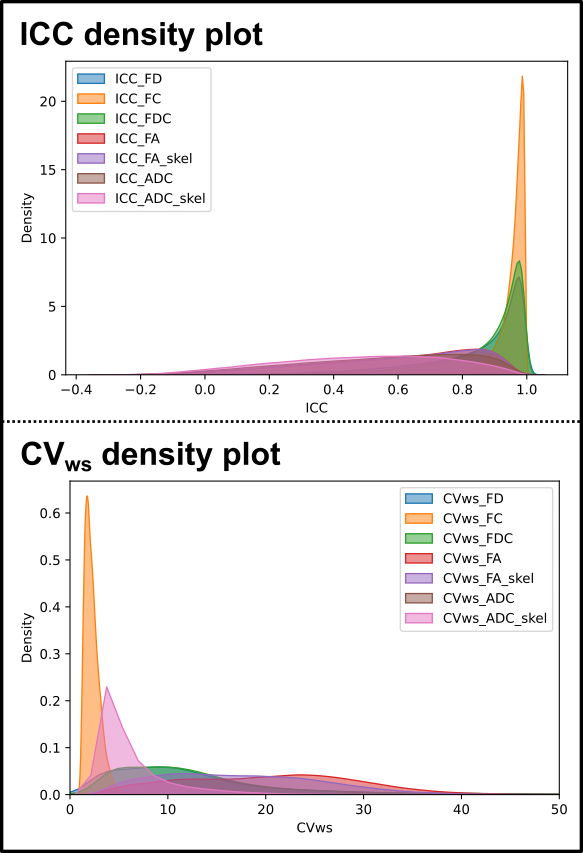

**Supplementary Figure S3: Density plots for ICC and CV_ws_ at each fixel/voxel.** The top plot shows distribution of ICC values across all fixels (for FBA metrics) and voxels (for DTI metrics) across the brain white matter. The bottom plot shows distribution density of CV_ws_ across brain white matter. For the FBA metrics, included fixels were defined on the FOD template image. For the DTI metrics, we used a voxel mask that corresponded to the fixel mask (including all voxels with containing white matter fixels). However, we additionally computed a constrained WM skeleton mask using an FA threshold of FA > 0.2. This data is summarised in Table 4.

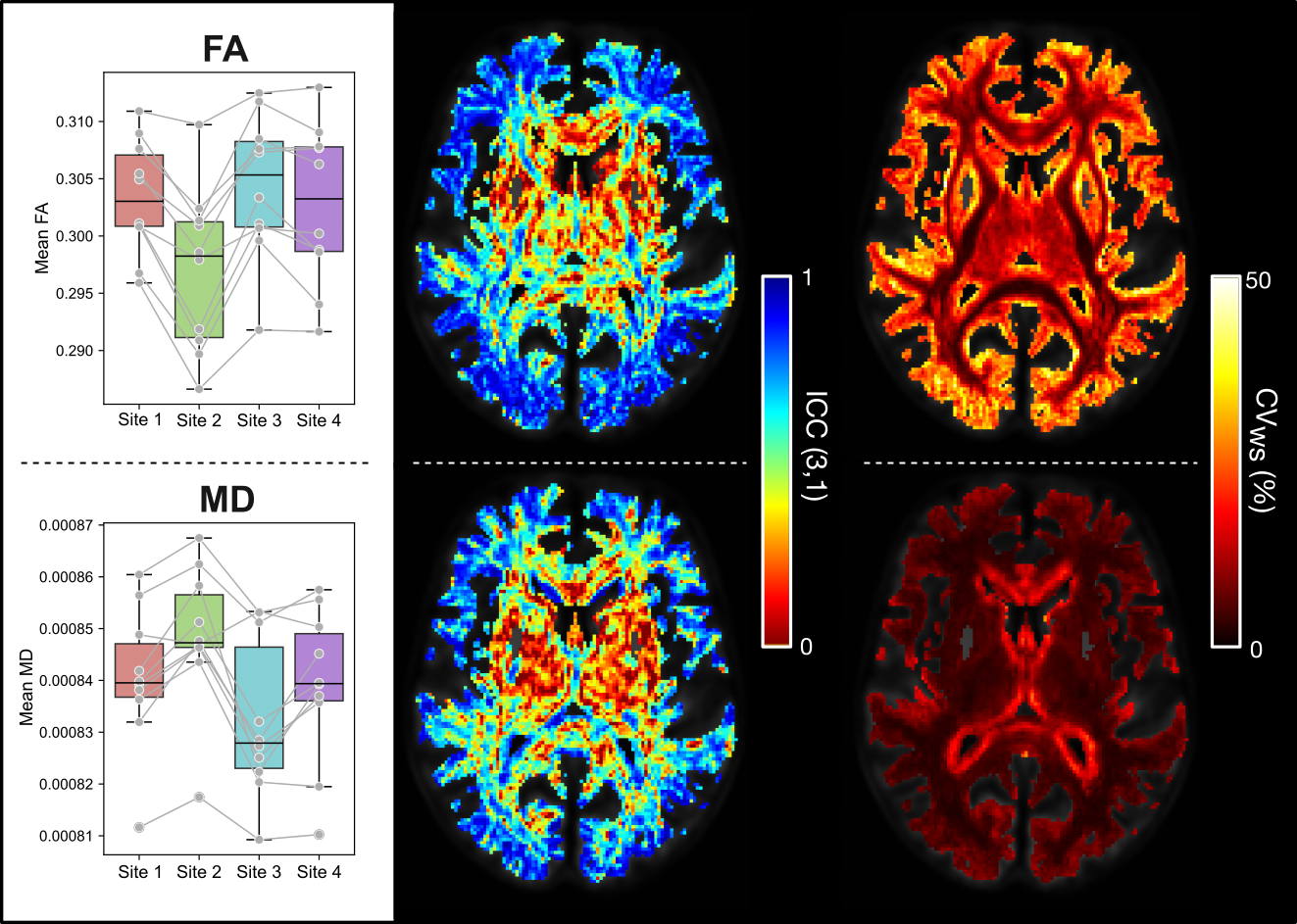

**Supplementary Figure S4: Reliability and reproducibility of tensor-based measures.** Top panel shows results for fractional anisotropy (FA), and bottom panel shows results for mean diffusivity (MD). Boxplots on the left show mean values across the whole brain, constrained to a white matter fixel mask (converted to a voxel mask). Middle images show ICC value at each voxel in the white matter voxel mask for a single axial slice. ICC values were generally lower in subcortical and cerebellar regions. Right images show CV_ws_ (%) at each voxel. The FA measure was generally more reproducible in key white matter regions, with higher variability towards cortical regions. MD had high reproducibility throughout the brain, with voxels surrounding the ventricles exhibiting slightly higher CV_ws_.

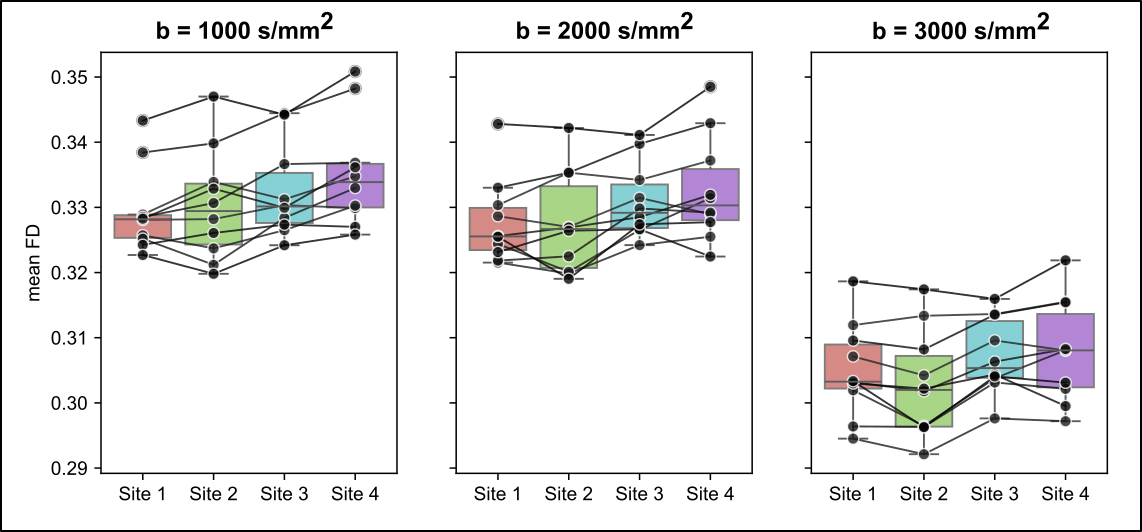

**Supplementary Figure S5: Boxplots showing mean FD across the whole brain at different *b*-values.** FD showed *b*-value dependence, but high and comparable reproducibility across all b-shells (see Supplementary Table S6).

**Table S1: CV and ICC of FD at the tract level (FD computed using site-specific response functions)**

| Tract | mean | | | std | | | CV_ws_ | CV_bs_ | | ICC(3,1) | | pval |
| --- | --- | --- | --- | --- | --- | --- | --- | --- | --- | --- | --- | --- |
| AF left | | 0.355 | | 0.014 | | | 1.11 | 3.96 | | 0.962 | | 0.0000000000 |
| AF right | | 0.355 | | 0.015 | | | 1.17 | 4.13 | | 0.966 | | 0.0000000000 |
| ATR left | | 0.325 | | 0.013 | | | 2.00 | 3.62 | | 0.865 | | 0.0000000000 |
| ATR right | | 0.330 | | 0.016 | | | 1.48 | 4.90 | | 0.958 | | 0.0000000000 |
| CA | | 0.342 | | 0.032 | | | 2.46 | 9.38 | | 0.964 | | 0.0000000000 |
| CC 1 | | 0.408 | | 0.024 | | | 3.63 | 5.29 | | 0.849 | | 0.0000000002 |
| CC 2 | | 0.390 | | 0.016 | | | 1.44 | 4.13 | | 0.951 | | 0.0000000000 |
| CC 3 | | 0.404 | | 0.025 | | | 2.20 | 6.23 | | 0.959 | | 0.0000000000 |
| CC 4 | | 0.512 | | 0.015 | | | 1.51 | 2.71 | | 0.912 | | 0.0000000000 |
| CC 5 | | 0.471 | | 0.016 | | | 1.48 | 3.26 | | 0.952 | | 0.0000000000 |
| CC 6 | | 0.434 | | 0.010 | | | 0.96 | 2.21 | | 0.935 | | 0.0000000000 |
| CC 7 | | 0.400 | | 0.015 | | | 0.74 | 3.84 | | 0.969 | | 0.0000000000 |
| CC | | 0.428 | | 0.012 | | | 1.13 | 2.79 | | 0.949 | | 0.0000000000 |
| CG left | | 0.322 | | 0.011 | | | 1.94 | 3.14 | | 0.894 | | 0.0000000000 |
| CG right | | | 0.313 | | 0.011 | 1.88 | | | 3.24 | | 0.889 | 0.0000000000 |
| CST left | | | 0.575 | | 0.015 | 1.43 | | | 2.38 | | 0.880 | 0.0000000000 |
| CST right | | | 0.584 | | 0.016 | 1.28 | | | 2.60 | | 0.912 | 0.0000000000 |
| FPT left | | | 0.481 | | 0.014 | 1.20 | | | 2.88 | | 0.925 | 0.0000000000 |
| FPT right | | | 0.488 | | 0.018 | 1.16 | | | 3.75 | | 0.966 | 0.0000000000 |
| FX left | | | 0.490 | | 0.032 | 1.65 | | | 6.63 | | 0.977 | 0.0000000000 |
| FX right | | | 0.451 | | 0.032 | 2.24 | | | 7.05 | | 0.980 | 0.0000000000 |
| ICP left | | | 0.359 | | 0.013 | 1.39 | | | 3.69 | | 0.915 | 0.0000000000 |
| ICP right | | | 0.338 | | 0.015 | 1.43 | | | 4.58 | | 0.937 | 0.0000000000 |
| IFO left | | | 0.378 | | 0.017 | 1.10 | | | 4.58 | | 0.954 | 0.0000000000 |
| IFO right | | | 0.362 | | 0.015 | 0.98 | | | 4.14 | | 0.960 | 0.0000000000 |
| ILF left | | | 0.367 | | 0.015 | 1.01 | | | 4.08 | | 0.947 | 0.0000000000 |
| ILF right | | | 0.357 | | 0.012 | 1.21 | | | 3.32 | | 0.946 | 0.0000000000 |
| MCP | | | 0.374 | | 0.015 | 1.17 | | | 4.01 | | 0.935 | 0.0000000000 |
| MLF left | | | 0.393 | | 0.011 | 0.91 | | | 2.92 | | 0.948 | 0.0000000000 |
| MLF right | | | 0.399 | | 0.012 | 1.29 | | | 2.78 | | 0.940 | 0.0000000000 |
| OR left | | | 0.381 | | 0.017 | 1.00 | | | 4.47 | | 0.966 | 0.0000000000 |
| OR right | | | 0.382 | | 0.014 | 0.98 | | | 3.82 | | 0.966 | 0.0000000000 |
| POPT left | | | 0.488 | | 0.018 | 1.20 | | | 3.63 | | 0.962 | 0.0000000000 |
| POPT right | | | 0.499 | | 0.015 | 1.24 | | | 2.92 | | 0.941 | 0.0000000000 |
| SCP left | | | 0.400 | | 0.014 | 1.01 | | | 3.43 | | 0.928 | 0.0000000000 |
| SCP right | | | 0.367 | | 0.016 | 1.12 | | | 4.54 | | 0.943 | 0.0000000000 |
| SLF III left | | | 0.353 | | 0.018 | 1.08 | | | 5.29 | | 0.975 | 0.0000000000 |
| SLF III right | | | 0.346 | | 0.010 | 1.25 | | | 2.75 | | 0.941 | 0.0000000000 |
| SLF II left | | | 0.337 | | 0.020 | 2.36 | | | 5.75 | | 0.943 | 0.0000000000 |
| SLF II right | | | 0.348 | | 0.014 | 1.54 | | | 3.79 | | 0.943 | 0.0000000000 |
| SLF I left | | | 0.325 | | 0.016 | 2.01 | | | 4.83 | | 0.949 | 0.0000000000 |
| SLF I right | | | 0.338 | | 0.015 | 2.02 | | | 4.12 | | 0.947 | 0.0000000000 |
| ST FO left | | | 0.318 | | 0.014 | 3.09 | | | 3.59 | | 0.652 | 0.0000065823 |
| ST FO right | | | 0.327 | | 0.015 | 2.89 | | | 3.82 | | 0.768 | 0.0000000401 |
| ST OCC left | | | 0.394 | | 0.017 | 0.98 | | | 4.45 | | 0.967 | 0.0000000000 |
| ST OCC right | | | 0.390 | | 0.015 | 1.04 | | | 3.83 | | 0.970 | 0.0000000000 |
| ST PAR left | | | 0.432 | | 0.015 | 1.04 | | | 3.48 | | 0.970 | 0.0000000000 |
| ST PAR right | | | 0.438 | | 0.011 | 1.24 | | | 2.25 | | 0.927 | 0.0000000000 |
| ST POSTC left | | | 0.444 | | 0.017 | 1.37 | | | 3.73 | | 0.970 | 0.0000000000 |
| ST POSTC right | | | 0.463 | | 0.012 | 1.37 | | | 2.38 | | 0.931 | 0.0000000000 |
| ST PREC left | | | 0.457 | | 0.013 | 1.31 | | | 2.76 | | 0.939 | 0.0000000000 |
| ST PREC right | | | 0.476 | | 0.012 | 1.28 | | | 2.28 | | 0.926 | 0.0000000000 |
| ST PREF left | | | 0.371 | | 0.013 | 1.39 | | | 3.33 | | 0.930 | 0.0000000000 |
| ST PREF right | | | 0.375 | | 0.017 | 1.27 | | | 4.64 | | 0.967 | 0.0000000000 |
| ST PREM left | | | 0.362 | | 0.019 | 1.72 | | | 5.21 | | 0.961 | 0.0000000000 |
| ST PREM right | | | 0.403 | | 0.025 | 1.90 | | | 6.16 | | 0.971 | 0.0000000000 |
| STR left | | | 0.577 | | 0.023 | 1.43 | | | 3.94 | | 0.965 | 0.0000000000 |
| STR right | | | 0.582 | | 0.021 | 1.07 | | | 3.62 | | 0.960 | 0.0000000000 |
| T OCC left | | | 0.380 | | 0.017 | 0.99 | | | 4.46 | | 0.968 | 0.0000000000 |
| T OCC right | | | 0.382 | | 0.014 | 0.96 | | | 3.59 | | 0.962 | 0.0000000000 |
| T PAR left | | | 0.427 | | 0.015 | 1.09 | | | 3.56 | | 0.968 | 0.0000000000 |
| T PAR right | | | 0.440 | | 0.013 | 1.23 | | | 2.82 | | 0.946 | 0.0000000000 |
| T POSTC left | | | 0.450 | | 0.018 | 1.37 | | | 3.89 | | 0.967 | 0.0000000000 |
| T POSTC right | | | 0.470 | | 0.012 | 1.31 | | | 2.27 | | 0.914 | 0.0000000000 |
| T PREC left | | | 0.479 | | 0.013 | 1.39 | | | 2.58 | | 0.917 | 0.0000000000 |
| T PREC right | | | 0.498 | | 0.015 | 1.26 | | | 2.81 | | 0.946 | 0.0000000000 |
| T PREF left | | | 0.389 | | 0.013 | 1.46 | | | 3.11 | | 0.921 | 0.0000000000 |
| T PREF right | | | 0.381 | | 0.018 | 1.28 | | | 4.66 | | 0.973 | 0.0000000000 |
| T PREM left | | | 0.372 | | 0.017 | 1.93 | | | 4.46 | | 0.939 | 0.0000000000 |
| T PREM right | | | 0.405 | | 0.018 | 2.04 | | | 4.23 | | 0.942 | 0.0000000000 |
| UF left | | | 0.346 | | 0.017 | 2.21 | | | 4.81 | | 0.930 | 0.0000000000 |
| UF right | | | 0.335 | | 0.019 | 2.12 | | | 5.49 | | 0.916 | 0.0000000000 |

**Table S2: CV and ICC of FC at the tract level (FC computed using site-specific response functions)**

| Tract | mean | std | CV_ws_ | CV_bs_ | ICC(3,1) | pval |
| --- | --- | --- | --- | --- | --- | --- |
| AF left | 1.046 | 0.080 | 0.80 | 7.91 | 0.994 | 0.0000000000 |
| AF right | 1.051 | 0.079 | 0.86 | 7.77 | 0.994 | 0.0000000000 |
| ATR left | 1.030 | 0.068 | 0.90 | 6.83 | 0.993 | 0.0000000000 |
| ATR right | 1.031 | 0.069 | 0.91 | 6.92 | 0.992 | 0.0000000000 |
| CA | 1.037 | 0.099 | 0.94 | 9.91 | 0.993 | 0.0000000000 |
| CC 1 | 1.065 | 0.103 | 1.53 | 9.93 | 0.984 | 0.0000000000 |
| CC 2 | 1.039 | 0.070 | 1.14 | 6.93 | 0.989 | 0.0000000000 |
| CC 3 | 1.033 | 0.077 | 1.49 | 7.65 | 0.981 | 0.0000000000 |
| CC 4 | 1.030 | 0.092 | 1.87 | 9.14 | 0.980 | 0.0000000000 |
| CC 5 | 1.013 | 0.069 | 1.80 | 6.95 | 0.966 | 0.0000000000 |
| CC 6 | 1.021 | 0.074 | 0.80 | 7.47 | 0.991 | 0.0000000000 |
| CC 7 | 1.121 | 0.116 | 0.84 | 10.74 | 0.998 | 0.0000000000 |
| CC | 1.033 | 0.070 | 0.98 | 7.01 | 0.988 | 0.0000000000 |
| CG left | 1.024 | 0.072 | 0.93 | 7.28 | 0.991 | 0.0000000000 |
| CG right | 1.023 | 0.078 | 1.11 | 7.84 | 0.991 | 0.0000000000 |
| CST left | 1.019 | 0.087 | 1.30 | 8.78 | 0.991 | 0.0000000000 |
| CST right | 1.024 | 0.080 | 1.37 | 8.07 | 0.990 | 0.0000000000 |
| FPT left | 1.018 | 0.069 | 0.93 | 7.03 | 0.993 | 0.0000000000 |
| FPT right | 1.023 | 0.072 | 0.94 | 7.27 | 0.995 | 0.0000000000 |
| FX left | 0.971 | 0.074 | 1.25 | 7.82 | 0.985 | 0.0000000000 |
| FX right | 0.978 | 0.087 | 1.00 | 9.20 | 0.992 | 0.0000000000 |
| ICP left | 1.050 | 0.057 | 0.75 | 5.60 | 0.984 | 0.0000000000 |
| ICP right | 1.053 | 0.058 | 0.99 | 5.62 | 0.971 | 0.0000000000 |
| IFO left | 1.103 | 0.103 | 0.73 | 9.67 | 0.998 | 0.0000000000 |
| IFO right | 1.106 | 0.102 | 0.59 | 9.63 | 0.998 | 0.0000000000 |
| ILF left | 1.119 | 0.117 | 0.79 | 10.87 | 0.998 | 0.0000000000 |
| ILF right | 1.121 | 0.118 | 0.83 | 10.90 | 0.998 | 0.0000000000 |
| MCP | 1.069 | 0.057 | 0.66 | 5.50 | 0.986 | 0.0000000000 |
| MLF left | 1.035 | 0.082 | 0.74 | 8.22 | 0.995 | 0.0000000000 |
| MLF right | 1.033 | 0.093 | 0.77 | 9.34 | 0.996 | 0.0000000000 |
| OR left | 1.111 | 0.121 | 0.75 | 11.27 | 0.998 | 0.0000000000 |
| OR right | 1.117 | 0.119 | 0.77 | 11.07 | 0.998 | 0.0000000000 |
| POPT left | 1.000 | 0.067 | 0.88 | 6.88 | 0.989 | 0.0000000000 |
| POPT right | 1.002 | 0.080 | 0.82 | 8.29 | 0.993 | 0.0000000000 |
| SCP left | 1.037 | 0.064 | 0.54 | 6.44 | 0.993 | 0.0000000000 |
| SCP right | 1.034 | 0.061 | 0.71 | 6.08 | 0.988 | 0.0000000000 |
| SLF III left | 1.045 | 0.081 | 0.90 | 7.98 | 0.992 | 0.0000000000 |
| SLF III right | 1.048 | 0.073 | 0.93 | 7.17 | 0.991 | 0.0000000000 |
| SLF II left | 1.038 | 0.084 | 1.18 | 8.39 | 0.989 | 0.0000000000 |
| SLF II right | 1.042 | 0.075 | 1.09 | 7.46 | 0.991 | 0.0000000000 |
| SLF I left | 1.023 | 0.087 | 1.63 | 8.77 | 0.980 | 0.0000000000 |
| SLF I right | 1.027 | 0.084 | 1.39 | 8.38 | 0.986 | 0.0000000000 |
| ST FO left | 1.052 | 0.086 | 0.83 | 8.45 | 0.992 | 0.0000000000 |
| ST FO right | 1.058 | 0.086 | 0.99 | 8.44 | 0.989 | 0.0000000000 |
| ST OCC left | 1.114 | 0.121 | 0.76 | 11.32 | 0.998 | 0.0000000000 |
| ST OCC right | 1.118 | 0.114 | 0.78 | 10.63 | 0.998 | 0.0000000000 |
| ST PAR left | 1.010 | 0.065 | 0.91 | 6.64 | 0.985 | 0.0000000000 |
| ST PAR right | 1.013 | 0.069 | 0.89 | 7.05 | 0.988 | 0.0000000000 |
| ST POSTC left | 1.000 | 0.062 | 1.37 | 6.34 | 0.975 | 0.0000000000 |
| ST POSTC right | 0.998 | 0.058 | 1.42 | 5.86 | 0.970 | 0.0000000000 |
| ST PREC left | 1.018 | 0.075 | 1.49 | 7.55 | 0.980 | 0.0000000000 |
| ST PREC right | 1.021 | 0.076 | 1.57 | 7.66 | 0.981 | 0.0000000000 |
| ST PREF left | 1.034 | 0.065 | 1.00 | 6.53 | 0.989 | 0.0000000000 |
| ST PREF right | 1.040 | 0.070 | 1.00 | 6.96 | 0.992 | 0.0000000000 |
| ST PREM left | 1.022 | 0.080 | 1.26 | 8.02 | 0.987 | 0.0000000000 |
| ST PREM right | 1.023 | 0.058 | 1.22 | 5.76 | 0.978 | 0.0000000000 |
| STR left | 1.027 | 0.093 | 1.82 | 9.32 | 0.981 | 0.0000000000 |
| STR right | 1.019 | 0.080 | 1.66 | 8.03 | 0.982 | 0.0000000000 |
| T OCC left | 1.108 | 0.119 | 0.75 | 11.13 | 0.998 | 0.0000000000 |
| T OCC right | 1.112 | 0.115 | 0.77 | 10.76 | 0.998 | 0.0000000000 |
| T PAR left | 1.009 | 0.065 | 0.93 | 6.61 | 0.985 | 0.0000000000 |
| T PAR right | 1.006 | 0.070 | 0.87 | 7.24 | 0.988 | 0.0000000000 |
| T POSTC left | 1.002 | 0.066 | 1.31 | 6.78 | 0.979 | 0.0000000000 |
| T POSTC right | 0.997 | 0.060 | 1.29 | 6.18 | 0.976 | 0.0000000000 |
| T PREC left | 1.019 | 0.081 | 1.50 | 8.20 | 0.982 | 0.0000000000 |
| T PREC right | 1.021 | 0.081 | 1.48 | 8.18 | 0.984 | 0.0000000000 |
| T PREF left | 1.026 | 0.065 | 1.07 | 6.51 | 0.986 | 0.0000000000 |
| T PREF right | 1.031 | 0.070 | 1.06 | 7.04 | 0.991 | 0.0000000000 |
| T PREM left | 1.018 | 0.078 | 1.21 | 7.90 | 0.987 | 0.0000000000 |
| T PREM right | 1.015 | 0.064 | 1.20 | 6.49 | 0.983 | 0.0000000000 |
| UF left | 1.067 | 0.116 | 1.01 | 11.32 | 0.997 | 0.0000000000 |
| UF right | 1.084 | 0.111 | 0.69 | 10.59 | 0.996 | 0.0000000000 |

**Table S3: CV and ICC of FDC at the tract level (FDC computed using site-specific response functions)**

| Tract | mean | | | std | | CV_ws_ | CV_bs_ | ICC(3,1) | pval |
| --- | --- | --- | --- | --- | --- | --- | --- | --- | --- |
| AF left | | 0.374 | 0.040 | | 1.57 | | 11.00 | 0.986 | 0.0000000000 |
| AF right | | 0.376 | 0.039 | | 1.59 | | 10.63 | 0.989 | 0.0000000000 |
| ATR left | | 0.337 | 0.028 | | 2.19 | | 8.41 | 0.965 | 0.0000000000 |
| ATR right | | 0.342 | 0.035 | | 1.78 | | 10.55 | 0.984 | 0.0000000000 |
| CA | | 0.356 | 0.065 | | 3.31 | | 18.62 | 0.981 | 0.0000000000 |
| CC 1 | | 0.431 | 0.058 | | 5.03 | | 13.17 | 0.940 | 0.0000000000 |
| CC 2 | | 0.406 | 0.037 | | 2.08 | | 9.29 | 0.976 | 0.0000000000 |
| CC 3 | | 0.420 | 0.045 | | 3.28 | | 10.77 | 0.963 | 0.0000000000 |
| CC 4 | | 0.526 | 0.054 | | 3.26 | | 10.25 | 0.963 | 0.0000000000 |
| CC 5 | | 0.477 | 0.042 | | 3.17 | | 8.67 | 0.955 | 0.0000000000 |
| CC 6 | | 0.445 | 0.038 | | 1.35 | | 8.72 | 0.985 | 0.0000000000 |
| CC 7 | | 0.443 | 0.049 | | 1.32 | | 11.50 | 0.993 | 0.0000000000 |
| CC | | 0.443 | 0.036 | | 1.85 | | 8.32 | 0.976 | 0.0000000000 |
| CG left | | 0.330 | 0.031 | | 2.26 | | 9.58 | 0.974 | 0.0000000000 |
| CG right | | 0.322 | 0.028 | | 2.30 | | 8.80 | 0.977 | 0.0000000000 |
| CST left | | 0.585 | 0.057 | | 2.72 | | 9.88 | 0.974 | 0.0000000000 |
| CST right | | 0.596 | 0.054 | | 2.63 | | 9.07 | 0.973 | 0.0000000000 |
| FPT left | | 0.489 | 0.042 | | 1.92 | | 8.68 | 0.975 | 0.0000000000 |
| FPT right | | 0.499 | 0.047 | | 1.73 | | 9.77 | 0.988 | 0.0000000000 |
| FX left | | 0.472 | 0.040 | | 2.18 | | 8.63 | 0.969 | 0.0000000000 |
| FX right | | 0.436 | 0.028 | | 2.84 | | 6.12 | 0.937 | 0.0000000000 |
| ICP left | | 0.375 | 0.028 | | 1.73 | | 7.48 | 0.965 | 0.0000000000 |
| ICP right | | 0.354 | 0.029 | | 1.95 | | 8.48 | 0.961 | 0.0000000000 |
| IFO left | | 0.412 | 0.047 | | 1.43 | | 11.72 | 0.988 | 0.0000000000 |
| IFO right | | 0.396 | 0.043 | | 1.29 | | 11.12 | 0.991 | 0.0000000000 |
| ILF left | | 0.409 | 0.053 | | 1.47 | | 13.52 | 0.993 | 0.0000000000 |
| ILF right | | 0.395 | 0.046 | | 1.88 | | 12.01 | 0.993 | 0.0000000000 |
| MCP | | 0.400 | 0.031 | | 1.54 | | 8.04 | 0.972 | 0.0000000000 |
| MLF left | | 0.407 | 0.038 | | 1.25 | | 9.65 | 0.988 | 0.0000000000 |
| MLF right | | 0.413 | 0.043 | | 1.71 | | 10.61 | 0.990 | 0.0000000000 |
| OR left | | 0.418 | 0.056 | | 1.48 | | 13.96 | 0.995 | 0.0000000000 |
| OR right | | 0.422 | 0.045 | | 1.53 | | 11.09 | 0.992 | 0.0000000000 |
| POPT left | | 0.490 | 0.044 | | 1.95 | | 9.19 | 0.981 | 0.0000000000 |
| POPT right | | 0.501 | 0.044 | | 1.96 | | 8.96 | 0.981 | 0.0000000000 |
| SCP left | | 0.411 | 0.038 | | 1.23 | | 9.52 | 0.985 | 0.0000000000 |
| SCP right | | 0.376 | 0.038 | | 1.51 | | 10.30 | 0.981 | 0.0000000000 |
| SLF III left | | 0.372 | 0.037 | | 1.55 | | 10.37 | 0.985 | 0.0000000000 |
| SLF III right | | 0.364 | 0.031 | | 1.81 | | 8.79 | 0.984 | 0.0000000000 |
| SLF II left | | 0.354 | 0.042 | | 3.04 | | 12.05 | 0.970 | 0.0000000000 |
| SLF II right | | 0.367 | 0.040 | | 2.26 | | 11.03 | 0.982 | 0.0000000000 |
| SLF I left | | 0.337 | 0.039 | | 3.20 | | 11.72 | 0.967 | 0.0000000000 |
| SLF I right | | 0.351 | 0.041 | | 3.06 | | 11.91 | 0.978 | 0.0000000000 |
| ST FO left | | 0.335 | 0.035 | | 3.53 | | 10.50 | 0.916 | 0.0000000000 |
| ST FO right | | 0.347 | 0.038 | | 3.73 | | 11.00 | 0.938 | 0.0000000000 |
| ST OCC left | | 0.431 | 0.055 | | 1.49 | | 13.21 | 0.994 | 0.0000000000 |
| ST OCC right | | 0.428 | 0.045 | | 1.62 | | 10.78 | 0.993 | 0.0000000000 |
| ST PAR left | | 0.437 | 0.037 | | 1.66 | | 8.59 | 0.983 | 0.0000000000 |
| ST PAR right | | 0.444 | 0.035 | | 1.95 | | 8.09 | 0.978 | 0.0000000000 |
| ST POSTC left | | 0.444 | 0.040 | | 2.68 | | 9.15 | 0.975 | 0.0000000000 |
| ST POSTC right | | 0.461 | 0.035 | | 2.85 | | 7.49 | 0.960 | 0.0000000000 |
| ST PREC left | | 0.465 | 0.043 | | 2.74 | | 9.21 | 0.969 | 0.0000000000 |
| ST PREC right | | 0.485 | 0.043 | | 2.84 | | 8.86 | 0.970 | 0.0000000000 |
| ST PREF left | | 0.384 | 0.030 | | 1.91 | | 8.00 | 0.969 | 0.0000000000 |
| ST PREF right | | 0.391 | 0.038 | | 1.82 | | 10.05 | 0.986 | 0.0000000000 |
| ST PREM left | | 0.374 | 0.039 | | 2.46 | | 10.62 | 0.976 | 0.0000000000 |
| ST PREM right | | 0.414 | 0.036 | | 2.76 | | 8.58 | 0.961 | 0.0000000000 |
| STR left | | 0.592 | 0.072 | | 3.39 | | 12.38 | 0.977 | 0.0000000000 |
| STR right | | 0.594 | 0.063 | | 2.83 | | 10.79 | 0.980 | 0.0000000000 |
| T OCC left | | 0.416 | 0.055 | | 1.47 | | 13.68 | 0.994 | 0.0000000000 |
| T OCC right | | 0.420 | 0.044 | | 1.50 | | 10.87 | 0.992 | 0.0000000000 |
| T PAR left | | 0.432 | 0.037 | | 1.73 | | 8.69 | 0.982 | 0.0000000000 |
| T PAR right | | 0.444 | 0.036 | | 1.90 | | 8.18 | 0.978 | 0.0000000000 |
| T POSTC left | | 0.452 | 0.043 | | 2.64 | | 9.55 | 0.976 | 0.0000000000 |
| T POSTC right | | 0.467 | 0.038 | | 2.65 | | 8.19 | 0.966 | 0.0000000000 |
| T PREC left | | 0.489 | 0.048 | | 2.84 | | 9.79 | 0.970 | 0.0000000000 |
| T PREC right | | 0.508 | 0.048 | | 2.76 | | 9.60 | 0.974 | 0.0000000000 |
| T PREF left | | 0.400 | 0.031 | | 2.06 | | 7.74 | 0.964 | 0.0000000000 |
| T PREF right | | 0.394 | 0.039 | | 1.87 | | 10.27 | 0.988 | 0.0000000000 |
| T PREM left | | 0.382 | 0.038 | | 2.63 | | 10.11 | 0.970 | 0.0000000000 |
| T PREM right | | 0.414 | 0.036 | | 2.82 | | 8.60 | 0.962 | 0.0000000000 |
| UF left | | 0.368 | 0.056 | | 2.81 | | 15.69 | 0.987 | 0.0000000000 |
| UF right | | 0.361 | 0.054 | | 2.51 | | 15.42 | 0.983 | 0.0000000000 |

**Table S4: Reproducibility of FA at the tract level**

| Tract | mean | std | CV_ws_ | CV_bs_ | ICC(3,1) | pval |
| --- | --- | --- | --- | --- | --- | --- |
| AF left | 0.347 | 0.011 | 1.99 | 2.91 | 0.889 | 0.0000000000 |
| AF right | 0.354 | 0.012 | 1.97 | 2.93 | 0.878 | 0.0000000000 |
| ATR left | 0.320 | 0.015 | 2.90 | 4.10 | 0.877 | 0.0000000000 |
| ATR right | 0.324 | 0.014 | 1.81 | 4.34 | 0.950 | 0.0000000000 |
| CA | 0.340 | 0.024 | 2.75 | 7.00 | 0.943 | 0.0000000000 |
| CC 1 | 0.434 | 0.021 | 1.94 | 4.82 | 0.880 | 0.0000000000 |
| CC 2 | 0.357 | 0.014 | 3.12 | 3.07 | 0.816 | 0.0000000021 |
| CC 3 | 0.364 | 0.021 | 4.58 | 4.23 | 0.804 | 0.0000000046 |
| CC 4 | 0.404 | 0.011 | 2.32 | 2.08 | 0.756 | 0.0000000744 |
| CC 5 | 0.394 | 0.012 | 1.76 | 2.60 | 0.859 | 0.0000000001 |
| CC 6 | 0.383 | 0.006 | 0.65 | 1.48 | 0.879 | 0.0000000000 |
| CC 7 | 0.355 | 0.008 | 0.77 | 2.33 | 0.937 | 0.0000000000 |
| CC | 0.382 | 0.009 | 1.75 | 2.01 | 0.856 | 0.0000000001 |
| CG left | 0.340 | 0.012 | 2.53 | 2.77 | 0.816 | 0.0000000020 |
| CG right | 0.339 | 0.010 | 2.56 | 2.09 | 0.727 | 0.0000003125 |
| CST left | 0.445 | 0.014 | 1.86 | 2.91 | 0.916 | 0.0000000000 |
| CST right | 0.443 | 0.016 | 2.01 | 3.27 | 0.910 | 0.0000000000 |
| FPT left | 0.410 | 0.016 | 2.84 | 3.20 | 0.889 | 0.0000000000 |
| FPT right | 0.415 | 0.018 | 2.68 | 3.84 | 0.932 | 0.0000000000 |
| FX left | 0.386 | 0.022 | 3.23 | 5.23 | 0.741 | 0.0000001660 |
| FX right | 0.358 | 0.023 | 2.80 | 6.20 | 0.866 | 0.0000000000 |
| ICP left | 0.331 | 0.007 | 1.49 | 1.83 | 0.570 | 0.0000876941 |
| ICP right | 0.323 | 0.007 | 1.34 | 1.81 | 0.593 | 0.0000452390 |
| IFO left | 0.348 | 0.010 | 1.09 | 2.91 | 0.931 | 0.0000000000 |
| IFO right | 0.337 | 0.009 | 0.79 | 2.71 | 0.943 | 0.0000000000 |
| ILF left | 0.355 | 0.013 | 1.31 | 3.73 | 0.935 | 0.0000000000 |
| ILF right | 0.347 | 0.012 | 0.92 | 3.54 | 0.945 | 0.0000000000 |
| MCP | 0.356 | 0.006 | 1.27 | 1.37 | 0.486 | 0.0007192794 |
| MLF left | 0.349 | 0.007 | 0.68 | 2.06 | 0.911 | 0.0000000000 |
| MLF right | 0.357 | 0.007 | 0.70 | 1.97 | 0.910 | 0.0000000000 |
| OR left | 0.356 | 0.010 | 1.02 | 2.87 | 0.941 | 0.0000000000 |
| OR right | 0.358 | 0.007 | 0.85 | 1.98 | 0.873 | 0.0000000000 |
| POPT left | 0.398 | 0.013 | 1.09 | 3.17 | 0.958 | 0.0000000000 |
| POPT right | 0.406 | 0.012 | 0.88 | 2.91 | 0.955 | 0.0000000000 |
| SCP left | 0.374 | 0.010 | 1.22 | 2.43 | 0.806 | 0.0000000039 |
| SCP right | 0.358 | 0.010 | 1.47 | 2.55 | 0.766 | 0.0000000456 |
| SLF III left | 0.362 | 0.013 | 2.10 | 3.23 | 0.874 | 0.0000000000 |
| SLF III right | 0.366 | 0.012 | 1.57 | 3.05 | 0.914 | 0.0000000000 |
| SLF II left | 0.337 | 0.016 | 3.29 | 3.77 | 0.858 | 0.0000000001 |
| SLF II right | 0.356 | 0.015 | 2.56 | 3.55 | 0.880 | 0.0000000000 |
| SLF I left | 0.354 | 0.013 | 3.06 | 2.55 | 0.791 | 0.0000000103 |
| SLF I right | 0.361 | 0.014 | 2.69 | 3.19 | 0.856 | 0.0000000001 |
| ST FO left | 0.339 | 0.015 | 1.89 | 4.17 | 0.843 | 0.0000000003 |
| ST FO right | 0.351 | 0.015 | 1.82 | 4.07 | 0.870 | 0.0000000000 |
| ST OCC left | 0.362 | 0.010 | 0.94 | 2.67 | 0.941 | 0.0000000000 |
| ST OCC right | 0.359 | 0.008 | 0.86 | 2.24 | 0.904 | 0.0000000000 |
| ST PAR left | 0.373 | 0.010 | 0.87 | 2.69 | 0.956 | 0.0000000000 |
| ST PAR right | 0.380 | 0.009 | 0.91 | 2.40 | 0.926 | 0.0000000000 |
| ST POSTC left | 0.382 | 0.015 | 1.53 | 3.79 | 0.959 | 0.0000000000 |
| ST POSTC right | 0.393 | 0.016 | 1.52 | 3.94 | 0.948 | 0.0000000000 |
| ST PREC left | 0.392 | 0.013 | 1.91 | 2.99 | 0.919 | 0.0000000000 |
| ST PREC right | 0.395 | 0.016 | 1.87 | 3.88 | 0.942 | 0.0000000000 |
| ST PREF left | 0.341 | 0.013 | 2.84 | 2.85 | 0.814 | 0.0000000023 |
| ST PREF right | 0.345 | 0.013 | 2.03 | 3.31 | 0.927 | 0.0000000000 |
| ST PREM left | 0.354 | 0.019 | 3.65 | 4.52 | 0.890 | 0.0000000000 |
| ST PREM right | 0.373 | 0.021 | 3.86 | 4.62 | 0.856 | 0.0000000001 |
| STR left | 0.431 | 0.015 | 2.02 | 3.08 | 0.868 | 0.0000000000 |
| STR right | 0.444 | 0.017 | 1.55 | 3.72 | 0.963 | 0.0000000000 |
| T OCC left | 0.355 | 0.010 | 1.00 | 2.88 | 0.939 | 0.0000000000 |
| T OCC right | 0.358 | 0.008 | 0.79 | 2.07 | 0.897 | 0.0000000000 |
| T PAR left | 0.370 | 0.010 | 0.89 | 2.61 | 0.949 | 0.0000000000 |
| T PAR right | 0.378 | 0.008 | 0.75 | 2.21 | 0.935 | 0.0000000000 |
| T POSTC left | 0.384 | 0.014 | 1.40 | 3.49 | 0.940 | 0.0000000000 |
| T POSTC right | 0.401 | 0.013 | 1.13 | 3.27 | 0.941 | 0.0000000000 |
| T PREC left | 0.396 | 0.012 | 1.99 | 2.67 | 0.909 | 0.0000000000 |
| T PREC right | 0.402 | 0.015 | 1.95 | 3.52 | 0.938 | 0.0000000000 |
| T PREF left | 0.352 | 0.013 | 3.12 | 2.77 | 0.790 | 0.0000000108 |
| T PREF right | 0.348 | 0.014 | 2.38 | 3.54 | 0.932 | 0.0000000000 |
| T PREM left | 0.353 | 0.018 | 3.59 | 4.08 | 0.861 | 0.0000000001 |
| T PREM right | 0.371 | 0.019 | 3.82 | 4.10 | 0.865 | 0.0000000000 |
| UF left | 0.326 | 0.015 | 2.40 | 4.20 | 0.877 | 0.0000000000 |
| UF right | 0.320 | 0.014 | 1.89 | 4.18 | 0.880 | 0.0000000000 |

**Table S5: Reproducibility of MD at the tract level**

| Tract | mean | | | std | CV_ws_ | CV_bs_ | ICC(3,1) | pval |
| --- | --- | --- | --- | --- | --- | --- | --- | --- |
| AF left | | 7.37E-04 | 1.059E-05 | | 1.20 | 1.01 | 0.681 | 0.0000022976 |
| AF right | | 7.33E-04 | 1.226E-05 | | 1.01 | 1.48 | 0.900 | 0.0000000000 |
| ATR left | | 7.70E-04 | 1.321E-05 | | 1.64 | 0.98 | 0.501 | 0.0005174143 |
| ATR right | | 7.68E-04 | 1.612E-05 | | 1.37 | 1.79 | 0.840 | 0.0000000003 |
| CA | | 8.25E-04 | 2.628E-05 | | 1.97 | 2.79 | 0.883 | 0.0000000000 |
| CC 1 | | 7.98E-04 | 2.571E-05 | | 1.75 | 2.95 | 0.823 | 0.0000000012 |
| CC 2 | | 7.85E-04 | 1.545E-05 | | 1.53 | 1.50 | 0.775 | 0.0000000272 |
| CC 3 | | 7.96E-04 | 2.012E-05 | | 1.91 | 1.98 | 0.776 | 0.0000000252 |
| CC 4 | | 7.83E-04 | 1.889E-05 | | 1.63 | 2.03 | 0.761 | 0.0000000579 |
| CC 5 | | 8.10E-04 | 2.452E-05 | | 1.69 | 2.75 | 0.843 | 0.0000000003 |
| CC 6 | | 8.25E-04 | 1.872E-05 | | 0.93 | 2.20 | 0.879 | 0.0000000000 |
| CC 7 | | 8.58E-04 | 2.212E-05 | | 1.06 | 2.50 | 0.941 | 0.0000000000 |
| CC | | 7.98E-04 | 1.450E-05 | | 1.15 | 1.57 | 0.830 | 0.0000000007 |
| CG left | | 7.56E-04 | 1.553E-05 | | 1.42 | 1.69 | 0.812 | 0.0000000026 |
| CG right | | 7.54E-04 | 1.532E-05 | | 1.47 | 1.62 | 0.803 | 0.0000000050 |
| CST left | | 7.45E-04 | 1.589E-05 | | 1.57 | 1.69 | 0.759 | 0.0000000637 |
| CST right | | 7.46E-04 | 1.512E-05 | | 1.69 | 1.43 | 0.726 | 0.0000003302 |
| FPT left | | 7.80E-04 | 1.654E-05 | | 1.67 | 1.59 | 0.742 | 0.0000001530 |
| FPT right | | 7.78E-04 | 1.878E-05 | | 1.90 | 1.82 | 0.782 | 0.0000000177 |
| FX left | | 1.54E-03 | 1.425E-04 | | 1.98 | 9.44 | 0.963 | 0.0000000000 |
| FX right | | 1.47E-03 | 1.405E-04 | | 1.76 | 9.82 | 0.969 | 0.0000000000 |
| ICP left | | 7.31E-04 | 2.072E-05 | | 1.72 | 2.50 | 0.712 | 0.0000006242 |
| ICP right | | 7.31E-04 | 1.871E-05 | | 1.78 | 2.10 | 0.707 | 0.0000007621 |
| IFO left | | 7.93E-04 | 1.427E-05 | | 1.10 | 1.58 | 0.885 | 0.0000000000 |
| IFO right | | 7.95E-04 | 1.948E-05 | | 1.07 | 2.36 | 0.954 | 0.0000000000 |
| ILF left | | 8.11E-04 | 1.801E-05 | | 1.18 | 2.05 | 0.876 | 0.0000000000 |
| ILF right | | 8.08E-04 | 2.206E-05 | | 1.09 | 2.66 | 0.953 | 0.0000000000 |
| MCP | | 7.56E-04 | 1.853E-05 | | 1.72 | 2.00 | 0.711 | 0.0000006399 |
| MLF left | | 7.94E-04 | 1.676E-05 | | 1.01 | 2.00 | 0.839 | 0.0000000003 |
| MLF right | | 7.92E-04 | 1.789E-05 | | 0.90 | 2.20 | 0.886 | 0.0000000000 |
| OR left | | 7.96E-04 | 1.575E-05 | | 1.21 | 1.73 | 0.876 | 0.0000000000 |
| OR right | | 8.07E-04 | 2.227E-05 | | 1.30 | 2.62 | 0.907 | 0.0000000000 |
| POPT left | | 8.08E-04 | 1.757E-05 | | 1.29 | 1.93 | 0.781 | 0.0000000189 |
| POPT right | | 8.10E-04 | 1.951E-05 | | 1.31 | 2.21 | 0.817 | 0.0000000018 |
| SCP left | | 7.42E-04 | 1.820E-05 | | 1.66 | 2.05 | 0.776 | 0.0000000251 |
| SCP right | | 7.54E-04 | 1.939E-05 | | 1.97 | 1.98 | 0.812 | 0.0000000026 |
| SLF III left | | 7.35E-04 | 1.074E-05 | | 1.36 | 0.87 | 0.541 | 0.0001920842 |
| SLF III right | | 7.31E-04 | 1.228E-05 | | 1.22 | 1.35 | 0.863 | 0.0000000000 |
| SLF II left | | 7.51E-04 | 1.135E-05 | | 1.40 | 0.91 | 0.523 | 0.0003015519 |
| SLF II right | | 7.43E-04 | 1.425E-05 | | 1.32 | 1.59 | 0.805 | 0.0000000043 |
| SLF I left | | 7.72E-04 | 2.066E-05 | | 1.77 | 2.27 | 0.768 | 0.0000000390 |
| SLF I right | | 7.56E-04 | 1.756E-05 | | 1.65 | 1.89 | 0.757 | 0.0000000718 |
| ST FO left | | 7.53E-04 | 1.466E-05 | | 1.32 | 1.63 | 0.760 | 0.0000000626 |
| ST FO right | | 7.48E-04 | 1.831E-05 | | 1.40 | 2.21 | 0.882 | 0.0000000000 |
| ST OCC left | | 8.01E-04 | 1.594E-05 | | 1.21 | 1.75 | 0.865 | 0.0000000000 |
| ST OCC right | | 8.05E-04 | 2.338E-05 | | 1.20 | 2.82 | 0.944 | 0.0000000000 |
| ST PAR left | | 7.88E-04 | 1.418E-05 | | 1.11 | 1.58 | 0.741 | 0.0000001654 |
| ST PAR right | | 7.84E-04 | 1.718E-05 | | 1.09 | 2.06 | 0.831 | 0.0000000006 |
| ST POSTC left | | 7.54E-04 | 1.373E-05 | | 1.33 | 1.46 | 0.714 | 0.0000005607 |
| ST POSTC right | | 7.42E-04 | 1.582E-05 | | 1.43 | 1.79 | 0.784 | 0.0000000161 |
| ST PREC left | | 7.20E-04 | 1.253E-05 | | 1.45 | 1.23 | 0.661 | 0.0000047961 |
| ST PREC right | | 7.14E-04 | 1.315E-05 | | 1.46 | 1.37 | 0.715 | 0.0000005486 |
| ST PREF left | | 7.46E-04 | 1.225E-05 | | 1.50 | 1.02 | 0.649 | 0.0000074170 |
| ST PREF right | | 7.44E-04 | 1.482E-05 | | 1.32 | 1.69 | 0.886 | 0.0000000000 |
| ST PREM left | | 7.31E-04 | 1.311E-05 | | 1.70 | 1.03 | 0.643 | 0.0000092835 |
| ST PREM right | | 7.12E-04 | 1.353E-05 | | 1.50 | 1.42 | 0.834 | 0.0000000005 |
| STR left | | 6.94E-04 | 1.305E-05 | | 1.74 | 1.14 | 0.465 | 0.0011510147 |
| STR right | | 6.96E-04 | 1.532E-05 | | 1.92 | 1.47 | 0.593 | 0.0000452730 |
| T OCC left | | 7.99E-04 | 1.524E-05 | | 1.20 | 1.65 | 0.866 | 0.0000000000 |
| T OCC right | | 8.11E-04 | 2.266E-05 | | 1.26 | 2.67 | 0.914 | 0.0000000000 |
| T PAR left | | 7.93E-04 | 1.414E-05 | | 1.13 | 1.54 | 0.723 | 0.0000003787 |
| T PAR right | | 8.00E-04 | 1.986E-05 | | 1.22 | 2.33 | 0.821 | 0.0000000014 |
| T POSTC left | | 7.60E-04 | 1.519E-05 | | 1.42 | 1.63 | 0.767 | 0.0000000428 |
| T POSTC right | | 7.51E-04 | 1.908E-05 | | 1.66 | 2.17 | 0.798 | 0.0000000067 |
| T PREC left | | 7.19E-04 | 1.259E-05 | | 1.56 | 1.14 | 0.617 | 0.0000218350 |
| T PREC right | | 7.14E-04 | 1.312E-05 | | 1.62 | 1.21 | 0.613 | 0.0000248692 |
| T PREF left | | 7.52E-04 | 1.234E-05 | | 1.61 | 0.87 | 0.492 | 0.0006394216 |
| T PREF right | | 7.51E-04 | 1.499E-05 | | 1.52 | 1.54 | 0.786 | 0.0000000143 |
| T PREM left | | 7.36E-04 | 1.405E-05 | | 1.80 | 1.12 | 0.607 | 0.0000297241 |
| T PREM right | | 7.18E-04 | 1.452E-05 | | 1.88 | 1.22 | 0.624 | 0.0000171591 |
| UF left | | 7.87E-04 | 1.834E-05 | | 1.47 | 2.02 | 0.918 | 0.0000000000 |
| UF right | | 7.94E-04 | 1.940E-05 | | 1.47 | 2.15 | 0.937 | 0.0000000000 |

**Table S6: Reproducibility and reliability of whole-brain fibre density (FD) at different b-values.**

| b-shell | Mean | SD | CV_ws_ (%) | CV_bs_ (%) | ICC (3,1) |
| --- | --- | --- | --- | --- | --- |
| b1000 | 0.332 | 0.0077 | 1.00 | 2.24 | 0.921 |
| b2000 | 0.330 | 0.0072 | 1.04 | 2.05 | 0.879 |
| b3000 | 0.306 | 0.0073 | 1.01 | 2.31 | 0.914 |
